# Ongoing Replication Stress Response and New Clonal T Cell Development Discriminate Between Liver and Lung Recurrence Sites and Patient Outcomes in Pancreatic Ductal Adenocarcinoma

**DOI:** 10.1101/2022.05.04.490552

**Authors:** Jason M. Link, Carl Pelz, Patrick J. Worth, Sydney Owen, Dove Keith, Ellen M. Langer, Alison Grossblatt-Wait, Allison L. Creason, Julian Egger, Hannah Holly, Isabel English, Kevin MacPherson, Motoyuki Tsuda, Jeremy Goecks, Emek Demir, Adel Kardosh, Charles D. Lopez, Brett C. Sheppard, Alex Guimaraes, Brian Brinkerhoff, Terry K. Morgan, Gordon Mills, Jonathan Brody, Rosalie C. Sears

## Abstract

**Background and Aims:** Metastatic pancreatic adenocarcinoma (mPDAC) is lethal, yet a subset of patients who have metastatic disease that spreads only to the lung have better outcomes. We identified unique transcriptomic and immune features that distinguish patients who develop metastases in the liver (liver cohort) versus those with lung-avid but liver-averse mPDAC (lung cohort).

**Methods:** We used clinical data from the Oregon Pancreas Tissue Registry to identify PDAC patients with liver and/or lung metastases. Gene expression and genomic alteration data from 290 PDAC tumors were used to identify features unique to patients from the liver and lung cohorts. In parallel, T cell receptor sequencing data from 289 patients were used to identify immune features unique to patients in the lung cohort.

**Results:** Lung cohort patients had better survival outcomes than liver cohort patients. Primary tumors from patients in the liver cohort expressed a novel gene signature associated with ongoing replication stress (RS) response predictive of poor patient outcome independent from known subtypes. In contrast, patients with tumors lacking the RS response signature survived longer, especially if their tumors had alterations in DNA damage repair genes. A subset of patients in the lung cohort demonstrated new T cell clonal development in their primary and metastatic tumors leading to diverse peripheral blood TCR repertoires.

**Conclusion:** Liver-avid metastatic PDAC is associated with an ongoing RS response, whereas tumors lacking the RS response with ongoing T cell clonal responses may have unique vulnerabilities allowing long-term survival in patients with lung-avid, liver-averse metastatic PDAC.

## Introduction

Patients diagnosed with pancreatic ductal adenocarcinoma (PDAC) face a dismal prognosis with a median overall survival of 1 year and an 11% chance of survival at 5 years; patients who present with metastatic disease (∼50%) fare even worse. Of the ∼20% of patients eligible for potentially curative resection of the primary tumor, only 30-35% survive >5 years. However, a subset of PDAC patients (∼10%) who develop lung metastases survive significantly longer than patients with metastatic spread to other sites^14–17^; in some cases, these patients survive >5 years with untreated, indolent lung metastases^18^. Patients with lung-restricted metastatic PDAC typically have slowly progressing disease, and may benefit from a metastatectomy^19^. In contrast, the common presentation of liver metastases or recurrent disease in the liver is frequently associated with poorer patient outcomes. Although not completely understood, at least two reasons may explain why liver metastases are common and so deadly. First, all venous outflow from the pancreas initially passes through the liver via the portal vein, which likely promotes tumor cell seeding in the liver. Second, the liver itself has been shown to be a nurturing tumor microenvironment for PDAC cells^20^, in part, suppressing tumor immunity^21^ and allowing PDAC liver metastases to flourish^22^.

Another factor contributing to mortality in PDAC patients with liver metastases is the greater likelihood that their tumors will be basal-like^23(p6),24^; a previously described subtype of PDAC associated with poor outcomes. Over the last decade, many studies have categorized PDAC tumors and cell lines into 2 to 6 mutually exclusive subtypes based on gene expression of tumors^5,25–27^, and the surrounding stroma containing fibroblasts and infiltrating leukocytes^28,29^. Two consensus subtypes reproducibly emerge from these studies^30^: the basal-like/quasi-mesenchymal/squamoid subtype and the classical/ductal/glandular subtype. These are confirmed by metabolomic^31^, proteomic^32^ and histologic^33^ subtyping, as well as expression of reproducibly identified, hallmark subtype-specific proteins^34^. Single cell analyses have confirmed the existence of the two consensus subtypes (basal-like and classical), and have additionally identified single PDAC tumor cells with a blend of the two subtypes^35,36^, suggesting a temporal progression from one subtype to the other, general plasticity between subtypes, or multiple overlapping subtype states. Although patients with classical-subtype tumors survive longer on average than patients with basal-like-subtype tumors, patients with either subtype still experience rapid progression and very poor outcomes. Therefore, other factors, superimposed on subtype, likely determine and drive the fate of patients with PDAC.

Many tumors of the basal-like subtype exhibit gene expression related to replication stress (RS)^37^, defined by stalled replication forks during DNA synthesis caused by premature entry into S phase, aberrant DNA replication, transcription/replication collisions, abnormal DNA damage checkpoints or altered DNA damage repair^38^. Failure to resolve RS leads to replication fork collapse, DNA damage, cell cycle arrest and ultimately senescence or cell death. Although aberrantly proliferating cancer cells are unavoidably plagued by RS, some malignant cells evolve response mechanisms to tolerate it, and their ability to survive the pro-mutagenic consequences of replication stress is key to their heterogeneity and aggressiveness. Moreover, RS also leads to cytosolic ssDNA, and consequently a cell intrinsic anti-viral response mediated by the stimulator of interferon genes (STING) that drives inhibition of proliferation and an interferon-mediated immune response meant to incapacitate the cells with cytosolic ssDNA. Thus, the strategies that malignant cells develop to respond to and survive RS (and not just RS *per se*) are critical to selection of drugs that target RS as well as the immune response provoked by RS.

Effective PDAC tumor immunity is rare, and both vaccines and immune checkpoint inhibitors (ICI) designed to generate or reinvigorate anti-tumor T cells have generally failed PDAC patients^39,40^ despite success in other solid malignancies. However, exceptional cases exist, demonstrating that adaptive tumor immunity does occur naturally and can be effective in some patients^41^. It remains unclear whether immunotherapy requires the right drug given in tandem to effectively treat PDAC patients and/or whether only a subset of patients will be candidates for therapies that modulate tumor immunity. Although microsatellite instability, mismatch repair deficiency, high tumor mutation burden, and PD-1/PDL-1 are potential biomarkers for immunotherapy efficacy^42–45^, there has been little progress identifying biomarkers of the immune response itself that inform treatment decisions for PDAC. Although some precision approaches to monotherapies and immunotherapies can moderately enhance outcomes for a fraction of PDAC patients^44^, future success will likely require a more complete understanding of metastatic location, cell intrinsic and microenvironment vulnerabilities, and the tumor immune landscape.

In this study, we generated and interrogated large datasets of tumor DNA and RNA sequencing as well as T cell receptor (TCR) sequencing from matched blood and tumor samples to evaluate the differences between liver-avid versus liver-averse, lung-avid PDAC. We develop a gene-expression signature specific to liver-avid versus liver averse, lung avid disease independent of the basal-like versus classical subtypes, that predicts poor patient outcome; and we identify evidence of ongoing replication stress response associated with this aggressive form of PDAC. Conversely, we show that primary tumors lacking the RS response signature are sensitive to alterations in DNA damage repair genes leading to long-term patient survival. Furthermore, we identify patients with lung-avid, liver-averse disease as a subset of patients with tumors lacking the RS response who exhibit ongoing T cell clonal development in their tumors and resultant high peripheral TCR diversity. Our findings herein identify a gene expression signature associated with RS response and identify tumors lacking this response as an exploitable weakness of PDAC that along with a specific type of functional adaptive tumor immunity can lead to improved outcomes for patients diagnosed with this devastating disease.

## Methods

### Tissue Acquisition and Patient Consent

Patient data, blood, and tissues were obtained with informed consent in accordance with the Declaration of Helsinki and were acquired through the Oregon Pancreas Tissue Registry under Oregon Health & Science University IRB protocol #3609.

### Clinical Data Collection

Clinical course timepoints, patient demographics, stage, grade, nodal involvement, resection margins, and angiolymphatic invasion were provided as deidentified data by the OHSU cancer registry with quality control data verification in a subset by a surgical pathologist (TM). For all patients in the liver and lung cohorts, we abstracted the site of all lesions proven to be metastatic by biopsy, described as likely metastatic by a radiologist, or that significantly increased in size during progression or decreased in size during treatment as long as a radiologist described the lesion as suspicious for or concerning for metastasis. Clinical imaging was reviewed by AG to validate patient assignments to the liver and lung cohorts. Time to recurrence was calculated from the earliest of either the recurrence date provided by the OHSU cancer registry, or the date of earliest lesion abstracted from CT reports.

### Specimen Processing

Primary and metastatic PDAC tumor specimens from consented patients at OHSU were processed by the OHSU department of pathology and preserved by standard formalin fixation and paraffin embedding leading to 3μm serial sections stained for hematoxylin & eosin.

### Histology Data

H&E stained formalin-fixed paraffin-embedded (FFPE) tissue sections from regions corresponding to those extracted for RNA-seq and somatic alteration analyses were independently appraised by two independent pathologists blinded to clinical data for the histologic features shown in Table 1. Overall diagnostic reproducibility kappa statistic was 0.89 [excellent defined as > 0.6]. Consensus diagnoses were used for metric comparisons.

**Table 1:**
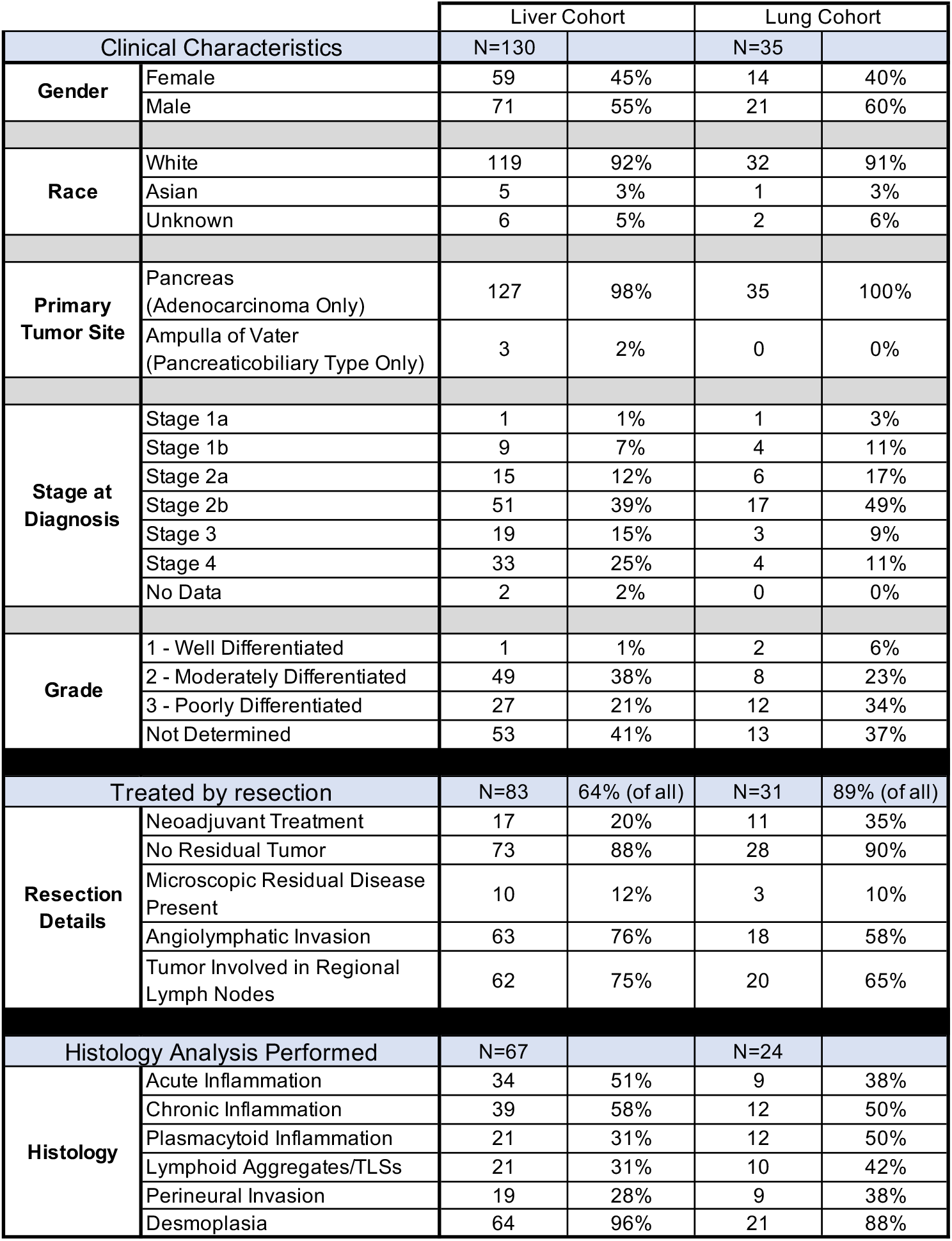
Patient demographics, disease characteristics, and tumor-specimen histology parameters for patients categorized into liver and lung cohorts. Percentages for resection details are only from primary tumor resections and percentages for histology are only from tumors with histology analyzed. Percentage of patients treated by resection was the only comparison significantly different between liver and lung cohorts (P<0.001, two-tailed Fisher’s exact test).

### Tempus RNA-seq and Genomic Alteration panel processing

OHSU provided FFPE PDAC specimen blocks along with matched normal blood or tissue to Tempus as part of a contract agreement. OHSU pathologist (TM) and Tempus pathologists marked regions of high tumor content (>20% ratio of tumor to normal nuclei) on H&E stained slides for DNA and RNA extraction. Solid tumor total nucleic acid was extracted from these tumor regions on adjacent FFPE tissue sections using Chemagic 360 sample-specific extraction kits (Perkin Elmer) and digested by proteinase K. RNA was purified from the total nucleic acid by DNase-I digestion. DNA sequencing of 596 genes and whole-transcriptome RNA sequencing were performed as previously described^1^.

Briefly, 100 nanograms (ng) of DNA for each tumor sample was mechanically sheared to an average size of 200 base pairs (bp) using a Covaris Ultrasonicator. DNA libraries were prepared using the KAPA Hyper Prep Kit, hybridized to the xT probe set, and amplified with the KAPA HiFi HotStart ReadyMix. One hundred ng of RNA for each tumor sample was heat fragmented in the presence of magnesium to an average size of 200 bp. Library preps were hybridized to the xGEN Exome Research Panel v1.0 (Integrated DNA Technologies) and target recovery was performed using Streptavidin-coated beads, followed by amplification with the KAPA HiFi Library Amplification Kit. The amplified target-captured DNA tumor library was sequenced using 2×126bp PE reads to an average unique on-target depth of 500x (tumor) and 150x (normal) on an Illumina HiSeq 4000. The amplified target-captured RNA tumor library was sequenced using 2×75bp PE reads to an average of 50M reads on an Illumina HiSeq 4000. Samples were further assessed for uniformity with each sample required to have 95% of all targeted bp sequenced to a minimum depth of 300x. Raw fastq files were returned to OHSU.

### DNA sequence analysis

DNA Variant detection, visualization, and reporting were performed as previously described^1^. Alignment and mapping was to GRCh37 using Novo align + BWA. CNVs derived from proprietary tumor/normal match analysis using CNAtools. Genomic variants are displayed on waterfall plots using GenVisR R package software^2^. Statistics are Fisher’s Exact test to look for significantly changed alteration rates between cohorts.

### RNA sequencing analysis

Paired-end fastq sequences were trimmed using trim-galore (ver 0.6.3) and default parameters. Pseudoalignment was performed with kallisto (ver 0.44.0) using genome assembly GRCh38.p5 and gencode (ver 24) annotation; default parameters were used other than the number of threads. The Bioconda package bioconductor-tximport (ver 1.12.1) was used to create gene level counts and abundances (TPMs). Quality checks were assessed with FastQC (ver 0.11.8) and MultiQC (ver 1.7). Quality checks, read trimming, pseudoalignment, and quantitation were performed via a reproducible snakemake pipeline, and all dependencies for these steps were deployed within the anaconda package management system^3,4^. We downloaded the “Pancreatic Adenocarcinoma (TCGA, PanCancer Atlas)” dataset from cbioportal.org.

### PurIST analysis

PurIST subtype calls and scores were generated using the PurIST method ^5^ applied to our RNA-Seq data. The PurIST authors provide instructions, R scripts, and gene pairs on GitHub (https://github.com/naimurashid/PurIST).

### Development of pORG and pSUB signatures

A two-factor analysis with DESeq2^6^ was performed on RNA-Seq counts from 76 primary samples. The two factors modeled were: liver cohort vs lung cohort and basal-like vs classical (from PurIST subtyping). For pORG, the most significant DE genes from the liver vs lung factor (p-value <0.01) were selected, then genes that co-occurred in the top half of the ranked genes from the basal vs classical factor were excluded, resulting in a list of 78 upregulated genes (only genes up in liver cohort were selected). For pSUB, the most significant DE genes from basal-like vs classical factor (p-value <0.01) were selected, then genes that co-occurred in the top half of the ranked genes from the liver vs lung factor were excluded, resulting in a list of 362 upregulated genes, of these, the 100 most significant were selected..

### GSEA and GSVA analyses

The GSVA tool^7^ was used to calculate relative pORG and pSUB gene set scores across all primaries, all metastases, and all tumors and identify top/bottom quartile cohorts. GSEA^8^ was run on clinical cohorts, pORG, pSUB, and PurIST cohorts using the MSigDB database v7.5.1 Hallmark gene set collection^9^. The 7 genes used for the IRDS signature were: STAT1, IFI44, IFIT3, OAS1, IFIT1, G1P2, and MX1

### VIPER analyses and Immune cell type enrichment

The transcriptional regulon enrichment analysis was performed using VIPER with the TCGA PAAD ARACNe-inferred network^10,11^. Gene expression data was normalized prior to running viper by median centering and scaling. VIPER regulon scores for all primaries were used for cohort comparisons.

Immune Cell Type enrichment analysis was scored using Gene Set Variation Analysis (GSVA) with gene sets defining 16 distinct immune cell types^7,12,13^ calculated for all primaries for cohort comparisons. GSVA was performed using log transformed gene expression data with a Gaussian kernel for estimation of the cumulative density function and a normalized enrichment statistic of the KS-like random walk.

### Software

R version 3.6.0 was used for GSVA and VIPER. R version 4.1.2 was used with R packages DESeq2, GSVA, msigdbr, gplots, and ggplot. GSEA was run in JAVA using the command line interface.

### TCRB Sequencing

Frozen leukocytes and 25 mm thick curls of formalin fixed, paraffin-embedded tumor were submitted to Adaptive Biotechnologies for human TCRB sequencing. Analyses were performed using the Immunoseq tool provided by Adaptive Biotechnologies. Samples with fewer than 100 productive templates were excluded from analyses. The Diversity Metrics Tool was used for richness and evenness metrics, and the Differential Abundance Tool was used to assess overlap between paired tumor and blood samples. The percentage of tumor-exclusive clones was calculated from a list of all rearrangements with ≥10 templates found in both samples combined. For shared, clonal sequences within cohorts, the top 50 rearrangements (by frequency in each sample) were compiled for all samples, and the Immunoseq Sequence Search Tool was used to identify all samples in the cohort that contained any of those CDR3b sequences at any frequency. Only the CDR3b amino sequences found in at least 25% of samples in the cohort were considered shared, clonal sequences.

### Statistics

All statistical tests were performed with GraphPad Prism. A Mantel-Cox test was used to compare all Kaplan-Meier survival and recurrence curves. ANOVA was used for plots with multiple comparisons and t-tests were used for plots with single comparisons. For non-Gaussian data (e.g., number of tumor-discrete TCR sequences) we used Kruskal-Wallis tests. Pearson and Spearman correlation coefficients were generated for continuous variables and non-parametric data, respectively

## Results

### PDAC patients who develop lung and not liver metastases have significantly better outcomes than patients who develop liver metastases

From a de-identified dataset of 1,873 patients diagnosed with and/or treated for PDAC at our institution between 2004 and 2020, we collected a complete set of disease-relevant computed tomography (CT) scans that were available for 234 (Supplemental Dataset 1). From these scans, we identified 35 patients who developed lung metastases but never developed liver metastases (referred to as the “lung cohort”) and 130 patients who developed liver metastases (referred to as the “liver cohort”). Consistent with previous reports, we observed that lung cohort patients fare significantly better by median overall survival (819 days) than liver cohort patients (468 days– Figure 1A). The median survival time for the liver cohort did not noticeably change if patients with both liver and lung metastases were excluded (450 days – Figure 1A). Median survival was also significantly different between patients in the lung and liver cohorts when comparing only patients treated by surgical resection of their primary tumors (876 days vs 549 days, respectively – Supplemental Figure 1A). For patients treated by primary tumor resection, median time to recurrence was significantly longer for patients in the lung cohort than patients in the liver cohort (303 days vs 166 days, respectively – Figure 1B), as was survival after resection (784 vs 498 days, respectively – Figure 1C), which positively correlated with time to recurrence for both cohorts (Supplemental Figure 1B) underscoring the role that recurrence plays in outcomes. Similarly, patients in the lung cohort survived longer after recurrence than patients in the liver cohort, though this difference was not significant (397 days vs 290 days, respectively, P=0.056 – Supplemental Figure 1C), and there was a significant correlation between median survival after recurrence and survival after resection for both cohorts (Supplemental Figure 1D). In contrast, there was no correlation between time to recurrence and survival time after recurrence (Supplemental Figure 1E) suggesting that there are different mechanisms of disease progression during these two clinical time periods.

**Figure 1:**
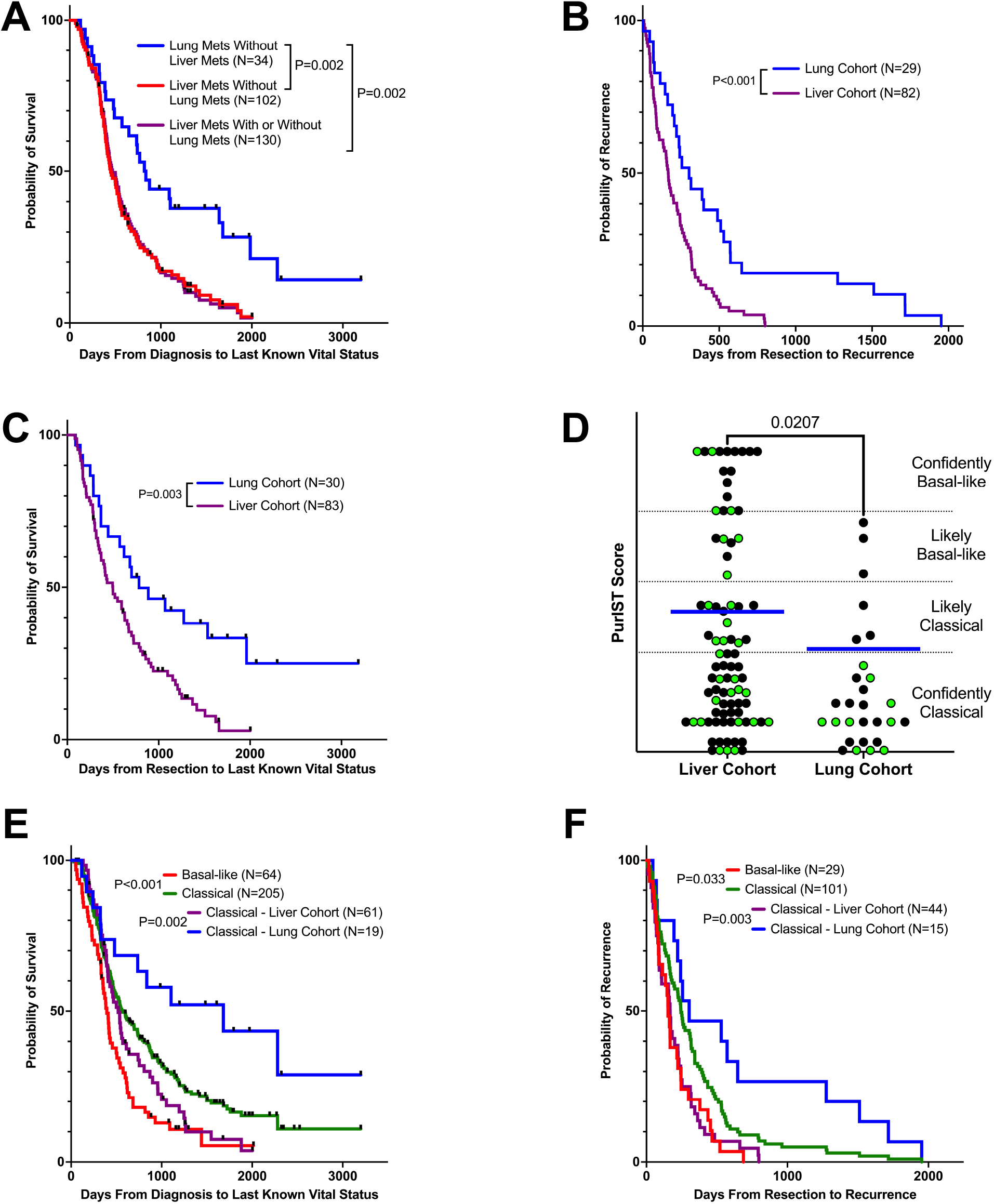
Outcomes for patients with liver metastases with or without lung metastases (liver cohort) and lung metastases without liver metastases (lung cohort): A) Overall survival, B) Days between resection and recurrence, and C) Survival since resection. D) PusIST subtyping scores for primary (black) and metastatic (green) tumor specimens from patients in the liver and lung cohorts. Blue bars represent means. Two-tailed Students t-test P value is reported. E) Overall survival and F) Days to recurrence for patients categorized by PurIST subtype and liver/lung cohorts. For charts with survival endpoints, patients not deceased were censored at last known vital status. Patients who died <30 days after resection were omitted. A Mantel-Cox test was used to compare all Kaplan-Meier survival and recurrence curves.

### The survival benefit from the classical subtype is not sufficient to explain the extended survival of lung cohort patients

We generated gene expression data by performing RNA-seq on histologically confirmed tumor-enriched regions from formalin-fixed paraffin embedded primary (N=218) and metastatic (N=72) PDAC tumors from patients at our institution, and then used PurIST^5^ to assign conventional subtypes of PDAC (basal-like or classical) to each tumor (Supplemental Dataset 1). Consistent with a previous report^24^, we found that tumors from patients in the lung cohort scored significantly more classical than basal-like (Figure 1D). Also consistent with previous reports^25,26^, we found an overall median survival benefit for patients with classical tumors compared to those with basal-like tumors (570 vs 394 days, respectively – Figure 1E). Median time to recurrence was also significantly longer for patients with classical-subtype tumors compared to those with basal-like tumors (244 vs 153 days, respectively – Figure 1F). However, when we assessed median survival of only patients with classical-subtype tumors in the lung vs liver cohorts, we found that patients in the lung cohort still lived significantly longer than patients in the liver cohort (1681 vs 520 days, respectively – Figure 1E) and also had a longer time to recurrence (303 vs 168 days, respectively – Figure 1F). These results indicate that there is a subtype-independent survival benefit for patients in the lung-cohort relative to the patients in the liver cohort.

### Clinical features and genomic alterations associated with metastatic-site avidity

We investigated clinical correlates of the liver and lung cohorts but did not observe differences in gender, race, stage at diagnosis, or tumor grade (Table 1). Patients in the lung cohort were more likely to be treated by resection than patients in the liver cohort (89% vs 64%, respectively – P<0.001), which may be due to more advanced disease at presentation for liver-cohort patients, however, the survival advantage in the lung cohort is still evident when only comparing patients treated by resection (Supplemental Fig 1A). A similar proportion of patients were treated with standard-of-care neoadjuvant chemotherapy in both cohorts and neoadjuvant treatment did not appear to influence time to recurrence for either cohort (Supplemental Figure 2).

Two surgical pathologists (TM and BB) performed a comprehensive histologic analysis of H&E stained serial sections adjacent to those extracted for RNA-seq. No significant histologic differences were noted between resection specimens from the two cohorts by grade or stage, including nodal involvement, presence of angiolymphatic invasion, presence of post-resection residual tumor, type of inflammation (acute versus chronic), or the presence/absence of prominent lymphoid aggregates (Table 1). Specimens from both cohorts showed extensive desmoplasia, as expected. We next determined the percentage of malignant cells in our specimens by using mutant allele frequencies determined from amplicon-based high-throughput sequencing of 595 genes known to be altered in cancer. We identified significantly higher tumor content in basal-like vs classical tumors (P=0.016 – Supplemental Figure 3); however, we did not identify a significant difference in the fraction of malignant cells in primary tumors between the liver and lung cohorts, or liver and lung metastases (Supplemental Figure 3).

We used the same gene panel DNA sequencing data to compare genes with alterations between basal-like and classical cohorts, and between liver and lung cohorts. The top 10 alterations are shown for each cohort (Supplemental Figure 2 and Supplemental Dataset 2). Among primary tumors, we found a significant increase in the frequency of alterations for ABL1, MKI67, and SMARCA4 in basal-like tumors, and for ATM, CUX1, GNAS, and TGFBR2 in classical tumors (Supplemental Table 1A). In liver-cohort primary tumors, alterations in TP53 were significantly more common, whereas PMS2 and PTEN alterations were more common in lung-cohort primary tumors (Supplemental Figure 4 and Supplemental Table 1A). Among metastases, we found 10 genes more likely to be altered in basal-like versus classical tumors (Supplemental Figure 4 and Supplemental Table 1B), but none that were different between the liver-cohort and lung-cohort metastases. Consistent with previous reports^46^, we found similar genes altered between paired tumors and metastases from the same patient (Supplemental Table 2).

### Gene-expression signatures in primary tumors specific to metastatic organotropism (pORG) or previously defined subtypes (pSUB) independently predict overall survival

Since outcome is influenced by recurrence site within the same subtype (Figure 1E and F), we sought to identify gene-expression pathways in primary tumors that may predict liver-avidity versus lung avidity/liver aversity of primary tumor cells without being influenced by the higher percentage of basal-like tumors in the liver cohort (Figure 1D). We ran a two-factor analysis with DESeq2^6^ to identify differentially expressed (DE) genes in primary tumors from the liver cohort vs lung cohort and from basal-like vs classical primary tumors (Supplemental Dataset 3). In order to focus on the biology of metastatic organotropism independent from previously published subtypes^25,28,30^, we excluded the top DE genes for established subtypes from the DE metastatic organotropism genes to obtain a <u>**p**</u>rimary <u>**org**</u>anotropism signature discriminating between liver-avid vs lung-avid, liver-averse tumors that we termed pORG (78 upregulated genes – Supplemental Dataset 3). We also performed this process for the DE <u>**p**</u>rimary tumor conventional <u>**sub**</u>type basal-like vs classical genes, subtracting the top DE organotropism genes to obtain a gene signature termed pSUB (100 upregulated genes -Supplemental Dataset 3).

We used the GSVA tool^7^ to run a Gene Set Variation Analysis of our primary tumor samples for both the pORG and pSUB gene sets. This analysis yields an activity score for each tumor relative to all other tumors (Supplemental Dataset 1). As expected, primary tumors from the liver cohort had significantly higher pORG scores than primary tumors from the lung cohort, but the pORG signature score did not significantly separate primary basal-like tumors from classical tumors (Figure 2A). Conversely, while pSUB significantly separated basal-like from classical tumors, it did not significantly separate the liver and lung cohort primary tumors (Figure 2A). We also produced pORG and pSUB GSVA scores for metastatic samples. These pORG scores did not distinguish liver-cohort from lung-cohort metastases (Figure 2B, left panel), indicating that this signature -and the functions it represents -are likely specific to primary tumors. Similar to other subtyping methods, the pSUB score did distinguish basal-like from classical metastases (Figure 2B, right panel) and also distinguished metastases from the liver and lung cohorts, underscoring previous reports that subtype biology carries over into metastases and that lung metastases are more likely to be classical (Figure 1D). GSVA scores for all specimens were not significantly different between all primary and all metastatic tumors for either pORG or pSUB (Supplemental Figure 5A).

**Figure 2:**
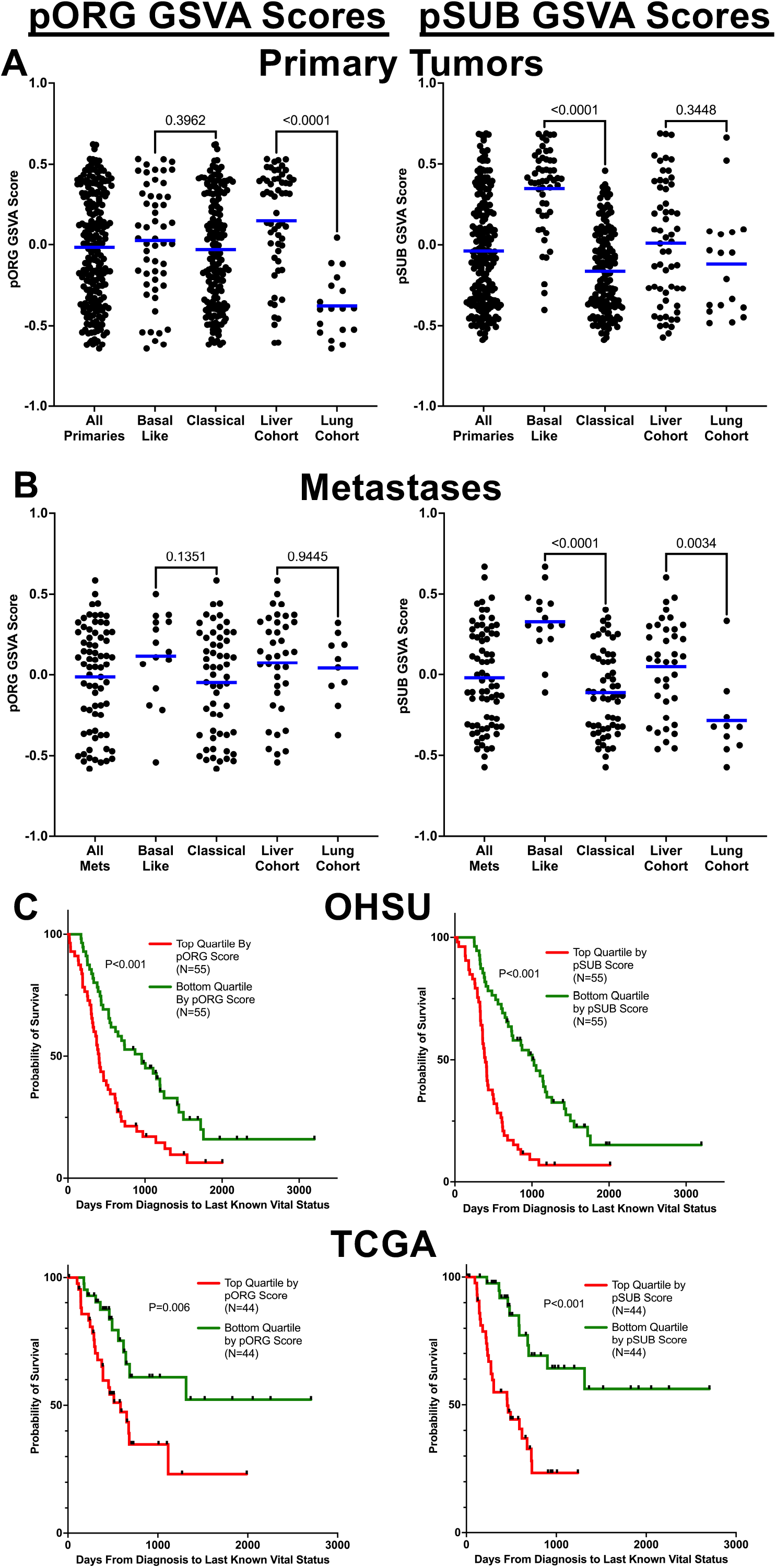
GSVA scores for the pORG (left panels) and pSUB (right panels) gene sets were calculated for the indicated cohorts from A) primary tumors and B) metastatic tumors. Blue bars represent means. One-way ANOVA was used to calculate indicated P values. C) Patient survival based on top and bottom quartiles of their primary tumors by pORG (left panels) and pSUB (right panels) GSVA scores from the OHSU dataset (top panels) and TCGA PAAD dataset (bottom panels). Patients who died within 30 days after resection are not shown. A Mantel-Cox test was used to compare Kaplan-Meier survival curves.

We found that overall survival significantly correlated with primary tumor pORG and pSUB GSVA scores independently (Supplemental Figure 5B). Relatedly, we found significant differences in overall survival between the top and bottom quartiles of all primary tumor specimens classified by pORG or pSUB GSVA scores for both our dataset (Figure 2C, top panels) and the pancreatic adenocarcinoma patient dataset (PAAD) reported by The Cancer Genome Atlas (Figure 2C, bottom panels). Thus, even though the pORG gene set is devoid of genes enriched in basal-like compared to classical tumors, and vice versa, each independently predicts outcome in our dataset and in the independent TCGA PAAD dataset, suggesting that each of these non-overlapping signatures can identify unique, clinically relevant biology.

### Distinct cancer hallmark pathways are enriched by pORG and pSUB gene sets

We used the cancer Hallmarks GSEA database to investigate pathways enriched in primary tumors with high vs low pORG scores and between the liver vs lung cohorts. We expected that specimens with high versus low pORG scores may more robustly identify GSEA pathways enriched in liver-avid primary tumors than the comparison between the liver and lung cohorts, as the latter comparison is influenced by subtype, and would be influenced by the inherent limitations of clinical-cohort assignments. Consistent with this, both high versus low pORG and liver-cohort versus lung-cohort comparisons showed similar pathway enrichments, but the liver vs lung comparison generally yielded lower normalized enrichment scores (Figure 3A and Supplemental Dataset 4). Both comparisons showed enrichment for pathways related to oncogene-mediated replication stress; specifically: G2M Checkpoint, E2F Targets, Mitotic Spindle, MYC Targets V1, and Interferon Alpha Response, as well as pathways related to cell metabolism and mitogenesis. The only hallmark pathway significantly enriched in low pORG or lung cohort was myogenesis. Specimens from both comparisons clustered by their most significantly enriched pathways (NES >1.7, nom p-value = <0.05) as expected (Figure 3B). We performed similar analyses of hallmark gene sets enriched in primary tumors comparing specimens defined as basal-like versus classical by PurIST, specimens with high versus low PurIST scores, and specimens with high versus low pSUB scores. All three of these comparisons revealed enrichment for pathways: Epithelial to Mesenchymal Transition (EMT), Apical Junction, hypoxia and angiogenesis, plausibly capturing the relationship between hypoxic stress and EMT^47^ (Figure 3C left panel and Supplemental Dataset 4). Additionally, the high pSUB score tumors showed enrichment of glycolysis and mTORC signaling. Inversely, we found pathways for Bile Acid Metabolism and Pancreas Beta Cells enriched in both classical subtype and low pSUB score primary tumors (Figure 3C and Supplemental Dataset 4). The significantly, positively enriched hallmark pathways (NES >1.7, nom p-value = <0.05) in these comparisons segregated tumor groupings as expected (Figure 3D). Together, these data indicate that separating metastatic organotropism and conventional molecular subtype gene expression by using our pORG and pSUB signatures can identify unique biologic pathway enrichments, with a high pORG score correlating with replication stress response pathways and a high pSUB score correlating with EMT and hypoxia pathways. Similar hallmark pathways were enriched in high pORG and high pSUB samples from the independent TCGA PAAD dataset (Supplemental Figure 6).

**Figure 3:**
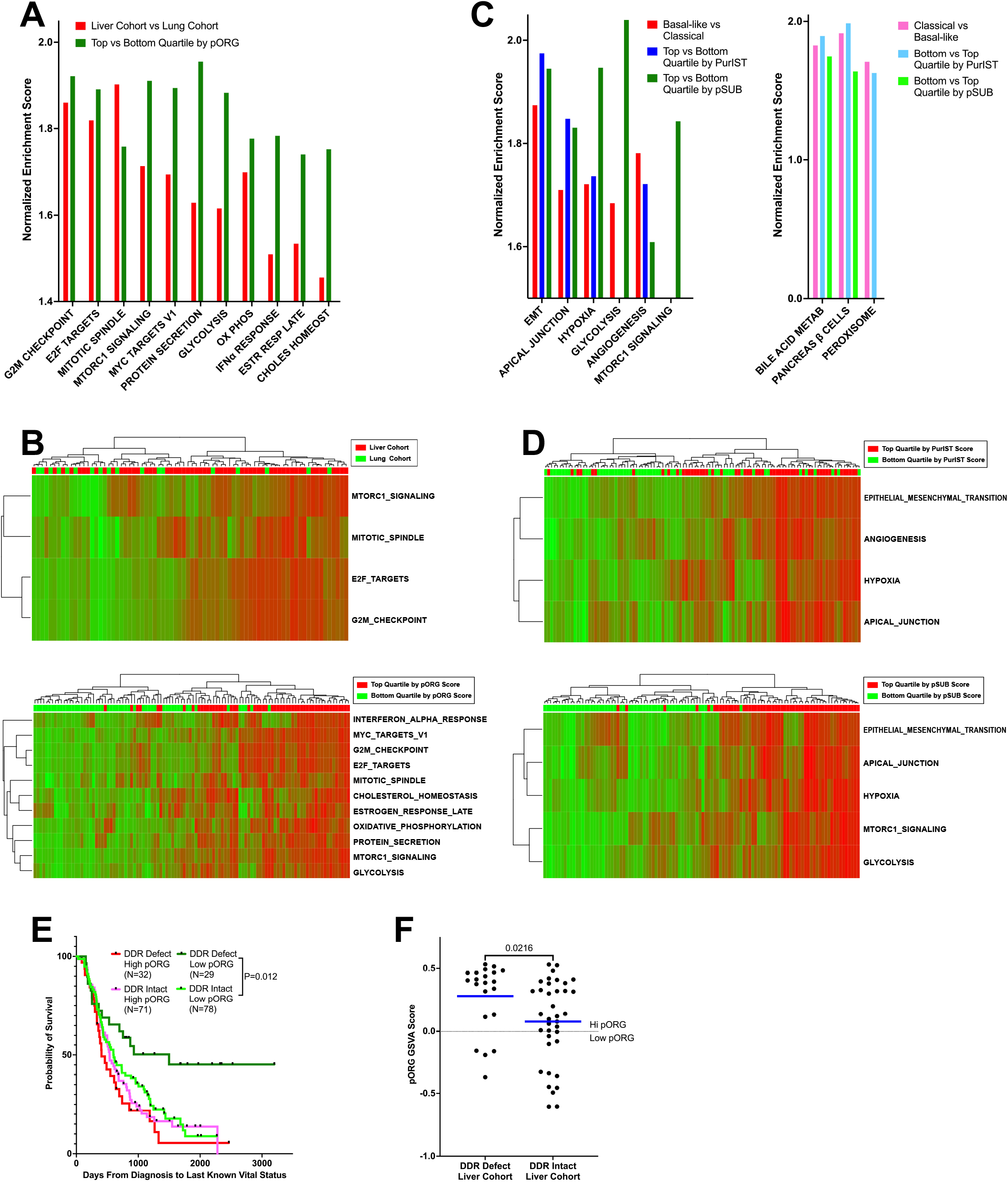
A) Normalized enrichment scores (NES) and are shown for hallmark GSEA pathways if any of the comparisons listed on each plot reached a NES >1.7 and nominal P<0.05. B) Heatmaps of GSVA scores (high=red, low=green) for primary tumors showing pathway (NES >1.7 and nominal P<0.05) and specimen clustering from the cohorts indicated on each plot. C) and D) Displayed as in A and B for the indicated cohort comparisons. E) Overall survival for patients with high or low pORG GSVA scores (top 50% or bottom 50%) stratified by known presence or absence of a somatic alteration in a DDR-related gene. A Mantel-Cox test was used to compare Kaplan-Meier survival curves. F) pORG GSVA scores for primary tumors from patients in the liver cohort categorized by a known alteration (DDR defect) or no variant detected (DDR intact) in a DDR-related gene. Two-tailed Students t-test P value is reported.

### Low pORG tumors are less tolerant to defects in DNA damage repair

A recent report by Dreyer et al. suggested that treatment-agent efficacy may depend on both replication stress (RS) and DNA Damage Response (DDR) gene alteration status – dividing patients into 4 categories based on the presence or absence of those two factors^37^. We considered that overall survival might be affected by these factors apart from treatment. Since our data indicate that liver-avid, high pORG score primary tumors are enriched for pathways indicative of ongoing RS, we divided patients into 4 similar categories by evenly splitting primary tumors by high and low pORG score and also by the presence or absence of a known DDR gene alteration^48^. Although patients with high pORG scoring tumors fared poorly regardless of DDR gene status, patients with low pORG scores survived significantly longer (P=0.012) if their primary tumors had altered DDR genes (Figure 3E). This result suggests that the effects of DDR gene alterations on tumor cell survival may be ameliorated in tumor cells with high pORG scores that have upregulated pathways responding to ongoing replication stress; consistent with activation of cell cycle checkpoint pathways to survive DNA damage and replication stress as well as MYC activity upregulating DNA damage response pathways in tumors with high pORG scores^49,50^. Additionally, liver-cohort primary tumors with DDR gene alterations had some of the highest pORG scores compared to those without DDR gene alterations (Figure 3F); suggesting that the presence of DDR gene alterations themselves may promote molecular mechanisms supporting tumor-cell responses to replication stress and DNA damage in order to avoid mitotic catastrophe.

### High pORG and liver-cohort tumors are enriched in transcription networks related to DNA damage repair, cell cycle promotion, and immune cell depletion

We performed transcriptional regulon enrichment analysis using VIPER and the TCGA pancreatic PAAD ARACNe-inferred network^10,11^ to identify regulatory protein activity scores between cohort comparisons (Supplemental Dataset 5). Consistent with GSEA results from the Hallmarks of Cancer gene sets (Figure 3), we observed significantly increased activity of proteins involved in DNA replication (ORC2 and MCM2-7 proteins – Figure 4A), and the response to replication stress (CHEK1, ATR, RAD51, BRCA1/2, BARD1, and PALB2 – Figure 4B) in high pORG vs low pORG and liver cohort versus lung cohort samples, while parallel proteins involved in double-strand break response like ATM and ARID1A were significantly decreased in activity (Figure 4B). Concurrently, positive cell cycle regulators (CyclinD, CDK2,4,6 and E2F1,2 – Figure 4C) were significantly increased in high versus low pORG and liver versus lung samples while the CDK inhibitor CDKN1B (P27) was significantly decreased (Figure 4C). We also found increased VIPER regulon activity scores in high vs low pORG for IFN alpha/beta receptor subunits 1 and 2 (mean DE 1.75 and 0.89, respectively – P<0.0001 for both), consistent with enrichment of IFNa response in high pORG samples by GSEA (Figure 3A and B). Together, these results support ongoing replication stress response mechanisms in high pORG/liver cohort PDAC.

**Figure 4:**
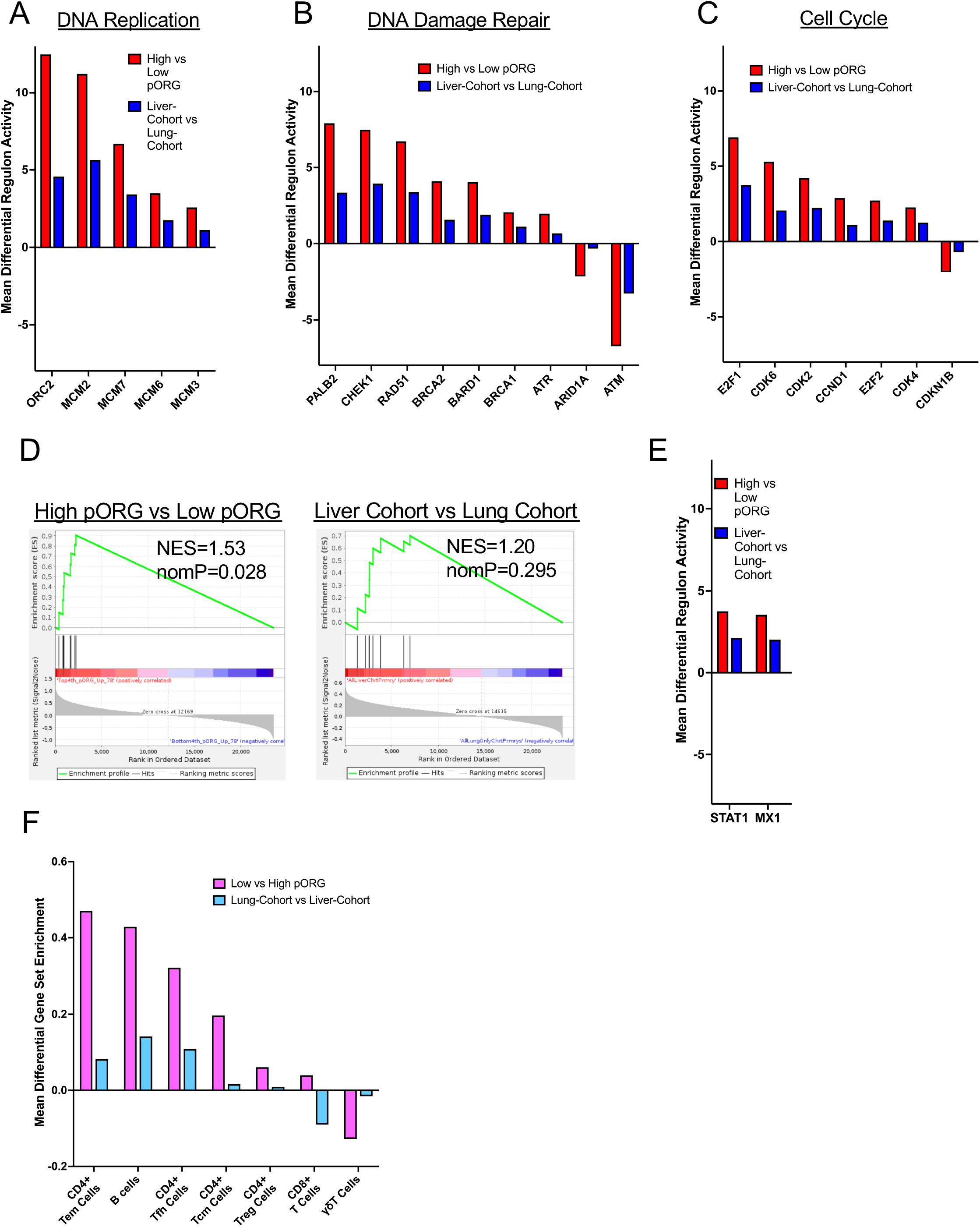
Mean differential VIPER regulon activity scores with significant (P<0.05 for all shown; one-way ANOVA) enrichment in high pORG vs low pORG (red) and liver-cohort vs lung-cohort (blue) primary tumors for regulons related to A) DNA replication B) DNA damage repair C) Cell cycle. D) GSEA plots and normalized enrichment scores (NES) for the IRDS gene set for the indicated comparisons. E) Mean differential VIPER regulon activity scores with significant (P<0.05 for all shown; one-way ANOVA) enrichment for 2 IRDS-related regulons in high pORG vs low pORG (red) and liver-cohort vs lung-cohort (blue) primary tumors. F) Enrichment of gene sets defining leukocyte subtypes in low pORG vs high pORG (pink) and lung-cohort vs liver-cohort (light blue) primary tumors (P<0.05 for pORG comparisons; one-way ANOVA).

The enrichment of gene expression related to Type 1 IFN (IFN) signaling in high pORG tumors does not match the well-known transient effects of IFN (inhibition of infected/tumor cell proliferation^51^ and upregulation of leukocyte effector functions^52^). However, the downstream activity of IFN is pleiotropic and changes dramatically over time; in fact, chronic IFN signaling in cancer induces an IFN-related DNA Damage Resistance Signature (IRDS) of gene expression that is associated with tumor cell resistance to DNA damage^53–55^ and escape from tumor immunity^56^. We used a set of IRDS genes reported to distinguish outcomes in cancer^53,55^ and found a significant enrichment in high pORG versus low pORG tumors (Figure 4D, left panel). A similar trend, though not significant, was seen in the liver-cohort versus lung-cohort primary tumors (Figure 4D, right panel). Two genes in the IRDS gene set matched VIPER regulons (STAT1 and MX1) and as expected, these were both significantly enriched in high pORG and liver-cohort tumors over low pORG and lung-cohort tumors, respectively (Figure 4E).

Chronic IFN signaling can also limit tumor immunity by favoring infiltration of suppressive leukocytes and inactivating adaptive immune cells^57^. To investigate this in our tumor cohorts we calculated the mean expression of sets of genes related to common lymphocyte subsets^12,58,59^. We found a significant enrichment of B cells and subsets of T cells (notably, CD4+ follicular helper and effector memory cells) in low pORG versus high pORG tumors (Figure 4F), and a similar pattern for lung-cohort versus liver-cohort tumors, though the latter comparisons were not significant. Taken together, these results suggest that tumors without ongoing RS and chronic IFN signaling are enriched for lymphocyte subtypes.

### Peripheral blood T cell receptor repertoires are more diverse in patients with lung-tropic, liver-averse disease

We performed sequencing of gene segments comprising the complementarity determining region 3 (CDR3) of T cell receptor□(TCRB) genes from 289 blood samples, 175 primary tumors (141 overlapping with the RNA-seq dataset), and 43 metastatic tumors (33 overlapping with the RNA-seq dataset). Within this dataset, we analyzed blood samples from 77 patients in the liver cohort and 16 patients in the lung cohort, of which 60 and 16 were matched with tumor samples from the same patient, respectively. Consistent with the gene-expression data showing enrichment for some T cell subsets in low pORG tumors (Figure 4F), we found a higher density of TCR templates in samples from low pORG tumors (Figure 5A, p= .0.012) but no difference between basal-like and classical subtypes. We did not find a higher density of TCR templates in lung-cohort tumors relative to liver-cohort tumors (Figure 5A), although this is consistent with the lack of a clear enrichment of T cells in lung-cohort over liver-cohort tumors (Figure 4F). As expected, the density of TCR templates per ng of tumor tissue significantly influenced the number of productive rearrangements we analyzed (i.e., the observed richness of the population, Supplemental Figure 7A), and we found significantly more productive rearrangements in low pORG tumors compared to high pORG tumors (Supplemental Figure 7B). These data suggest that mechanisms represented by a high pORG signature may restrict the density of T cells in the tumor and reduce the richness of analyzed TCR repertoires.

**Figure 5:**
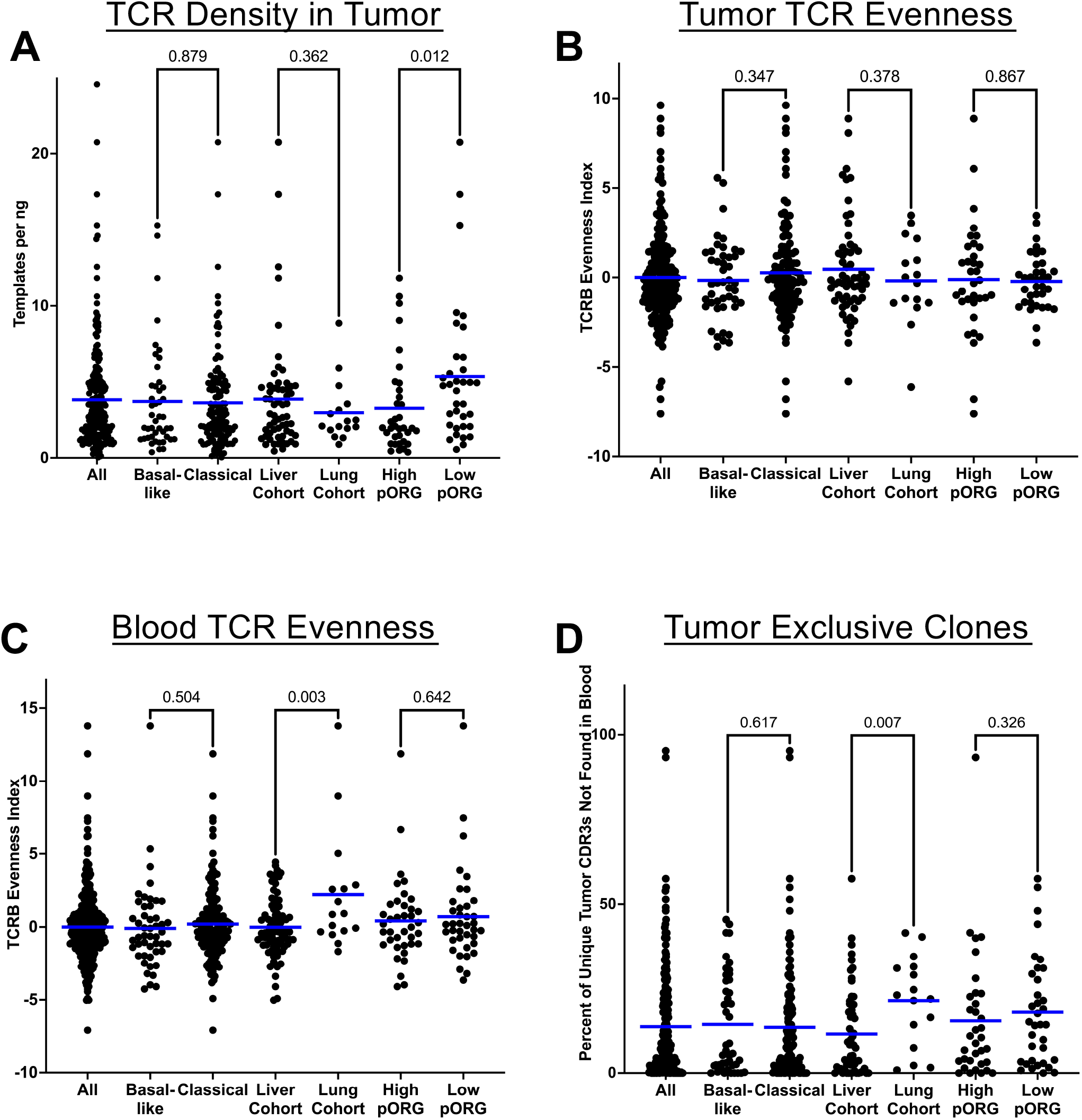
A) The density of productive TCRB templates (per ng of tumor tissue) in tumors from the indicated cohorts. B) The Evenness Index of the TCRB repertoires from tumors in each indicated cohort. C) The Evenness Index of the TCRB repertoires from blood samples in each indicated cohort. D) The percentage of unique tumor TCRB CDR3β sequences with ≥10 templates detected in tumor samples, but none detected in paired (i.e., from the same patient) blood samples. For all, blue bars represent means. One-way ANOVA was used to calculate indicated P values.

T cell repertoire diversity is a function of both richness (the abundance of TCRs) and evenness (the distribution of the frequencies of clones within a sample – a very clonal repertoire would have low evenness). We developed an Evenness Index that is the combination of three metrics unrelated to richness and unlikely to be affected by differences in the number of TCR productive rearrangements analyzed (Pielou Evenness, Simpsons Evenness, and Clone Distribution Slope)^60^. We did not find a difference in TCR repertoire evenness between any of the tumor cohorts indicating that the overall shapes of the liver-cohort and lung-cohort tumor repertoires are not different (Supplemental Figure 7C and Figure 5B). In contrast, we found significantly more TCR repertoire evenness (i.e., less clonality) within peripheral blood samples from patients in the lung cohort (Figure 5C). We considered that although evenness/clonality was similar between liver-cohort and lung-cohort tumors, that the nature of those clonal responses may differ, and that difference may manifest in the peripheral blood TCR. To investigate this, we used data from 214 matched pairs of tumor and blood samples (91% collected on the same day) and looked for unique TCRB clones in tumor samples that were absent in matched blood samples (i.e., tumor-distinct clones), suggestive of new clonal development. We found that lung-cohort tumors harbored significantly more of these tumor-distinct clones (Figure 5D). This result is consistent with ongoing, new T cell clonal development in lung-cohort tumors that is similar in evenness/clonality to other tumors at any one timepoint, but as new clones emigrate to the periphery, TCR diversity increases in blood. Consistent with this conclusion, we found a positive correlation between the Evenness Index in blood samples and the percentage of tumor-distinct clones for all 214 patients with matched blood and tumor samples (Supplemental Figure 7D). Additionally, we found that the Evenness Index in tumor samples negatively correlated with the percentage of tumor-distinct clones (Supplemental Figure 7E), underscoring that new clonal development contributes to overall clonality of the tumor T cell repertoire.

Our data suggest that T cell responses in liver-cohort tumors are not directed at new antigens over time, but instead may reflect chronic reactions to common PDAC-associated (or other) antigens present since tumor initiation. To investigate these responses, we selected published CDR3b sequences that have been experimentally confirmed as part of TCRs specific for KRAS G12/13 alterations^61^, a tumor-initiating mutation present in ∼90% of PDAC tumors. We identified 4 of these CDR3b sequences in liver cohort blood and tumor samples, and 3 of those 4 CDR3b sequences in lung-cohort blood and tumor samples (Supplemental Dataset 6). Consistent with repeated expansion of the same clones over time, we found these putative mutant KRAS-specific CDR3b sequences at significantly higher frequencies in the analyzed repertoires of liver-cohort tumor samples compared to the lung-cohort tumor samples (Figure 6A), suggesting that T cell responses to tumor-initiating neoepitopes are more likely to expand in liver-cohort tumors. These sequences existed at similarly low frequencies in liver-cohort and lung-cohort blood samples (Figure 6A), consistent with limited expansion of these clones in the periphery. The presence of these putative mutant KRAS-specific CDR3b sequences in tumors did not correlate with better outcomes; rather the median overall survival for patients with KRAS-specific CDR3β sequences in tumors was shorter for both liver-cohort (391 days) and lung-cohort (565 days) patients compared to the median survival for all patients in these cohorts (468 and 819 days, respectively – Figure 1A).

**Figure 6:**
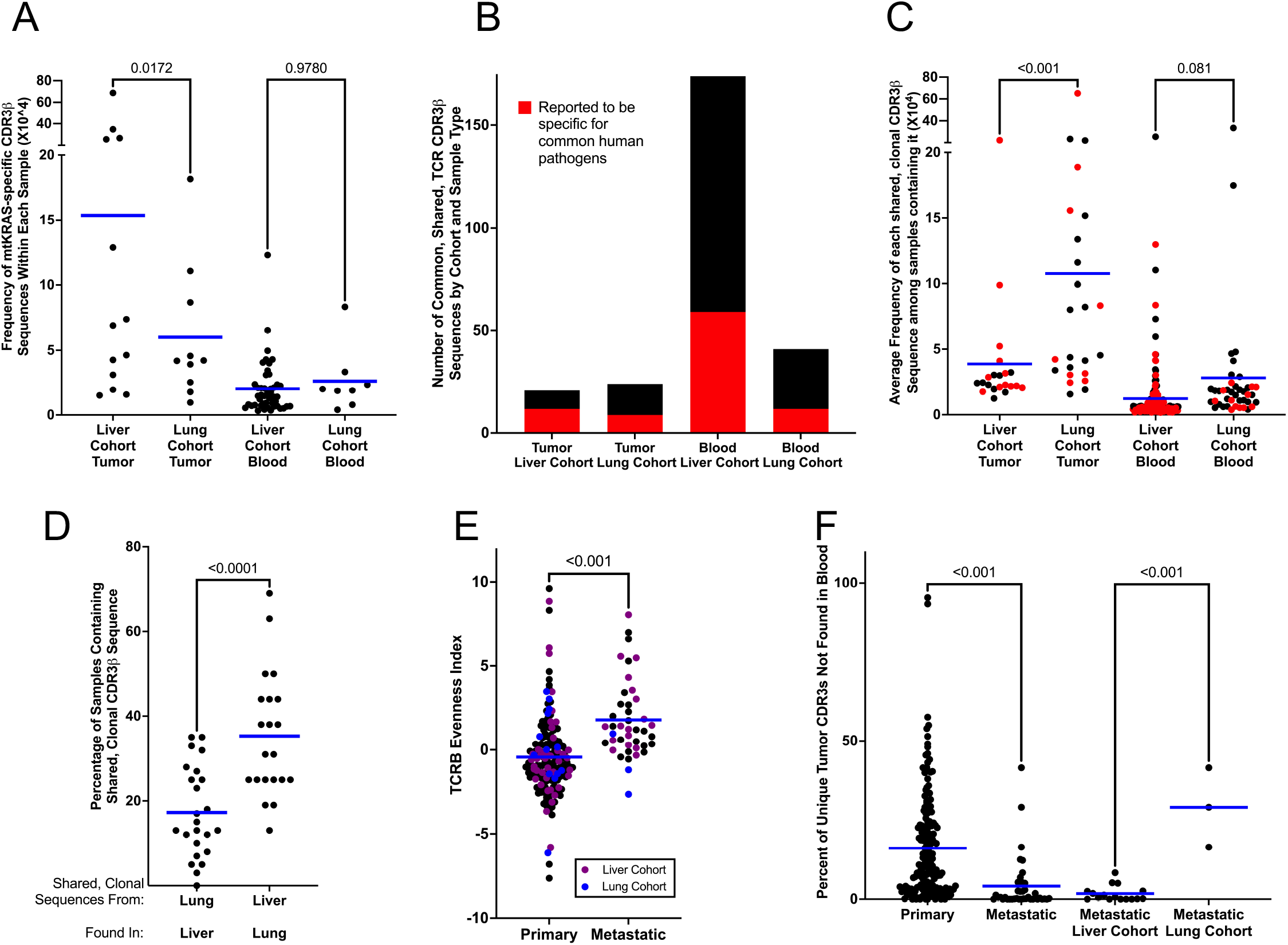
A) The frequency of any putative mutant KRAS-specific CDR3β sequences within the TCR repertoire for each sample containing at least one of those sequences (each dot represents one sample). B) The number of shared, clonal sequences found for each cohort; red fractions of bars represent the number of CDR3β sequences with reported specificity for common human pathogens. C) The average frequency of each shared, clonal CDR3β sequence in all samples containing that sequence (each dot represents one shared, clonal CDR3β sequence); red dots represent CDR3β sequences with reported specificity for a common human pathogen. D) The percentage of samples containing at least one of the alternate cohort’s shared, clonal sequences (each dot represents one CDR3β sequence). E) The Evenness Index score for all primary and metastatic samples; purple dots represent samples from liver-cohort patients and blue dots represent samples from lung-cohort patients. F) The percentage of tumor-distinct clones in primary and metastatic tumors. For all, blue bars represent means. One-way ANOVA was used to calculate indicated P values for A, C, and F. Two-tailed Student’s t-test was used to calculate P values for D and E.

Lung-cohort tumors are a subset of low-pORG tumors that may have capitalized on the higher density of tumor infiltrating T cells (Supplemental Fig 7E) to generate ongoing, multi-clonal responses in tumors. We considered that the number of public TCRs – i.e., identical TCRs specific for the same critically immunogenic epitopes – could represent common responses within each cohort. To investigate this, we identified the 50 most frequent CDR3β amino acid sequences from the TCR repertoire of each patient and compiled them in to four separate lists: liver-cohort blood, liver-cohort tumor, lung-cohort blood and lung-cohort tumor (Supplemental Dataset 6), and then looked for these clonal sequences in the entire TCR repertoires of all samples in the same cohort. We then identified any of these clonal CDR3b sequences that were shared by at least 25% of the samples in the same cohort. Thus, we identified CDR3b sequences that were *shared* by a large fraction of samples in the cohort and were dominantly *clonal* in at least one sample. None of the shared, clonal sequences matched those reported to be mutant KRAS specific^61^. We found similar numbers of these shared, clonal CDR3β sequences in liver-cohort tumors (N=21) as in lung-cohort tumors (N=24); in contrast, we found many more shared, clonal CDR3β sequences among blood samples from patients in the liver cohort (N=174) than from patients in the lung cohort (N=41, Figure 6B). This result is consistent with repeated responses to a limited set of common tumor-initiating neoepitopes in liver-cohort tumors over time, increasing the chance that tumors from different patients would generate identical TCRs. These shared, clonal sequences may accumulate in the periphery even though they are not found in many liver-cohort tumors at any one point in time. In contrast to responses to putative mutant KRAS-specific tumor-initiating epitopes, the shared, clonal responses in lung-cohort tumors, existed at higher frequencies than in liver-cohort tumors (Figure 6C). These results suggest that liver-cohort tumors may promote clonal responses against persistent neoepitopes (Figure 6A), whereas lung-cohort tumors may be more likely to make clonal responses against new (though uncharacterized) tumor antigens (Figure 6C) as they respond to new antigens over time (Figure 5D). As expected, some of the shared, clonal CDR3b sequences have been previously identified in TCRs known to recognize common infectious agents in humans^62^ (Figure 6B and C), suggesting that poor expansion of T cell clones may be a characteristic of liver-cohort patients regardless of relevance to tumor immunity. We considered that the shared, clonal sequences we identified in lung-cohort tumors may also be present in some liver-cohort tumors (and vice versa) as public TCRs are unlikely to be perfectly cohort-specific. In fact, we found the shared, clonal sequences from each tumor cohort in the other tumor cohort; but we were more likely to find shared, clonal liver-cohort tumor CDR3b sequences in lung-cohort tumors compared to the inverse (Figure 6D). This underscores that these public/shared, clonal sequences are being generated by both cohorts, but more commonly present (Figure 6D) and expanded (Figure 6C) in lung-cohort tumors.

PDAC tumors that promote new, clonal T cell responses over time may be more likely to generate T cell-mediated tumor immunity that suppresses growth of liver metastases. In lung-cohort patients, this seemingly more effective tumor immunity exists in patients concurrently with metastatic disease (even at resection, patients with future recurrent disease must have occult metastases), however, we expected that metastases would harbor less clonal outgrowth as progressive disease escapes tumor immunity. In fact, we found this to be true for metastases in general, as they had significantly greater evenness in their TCR repertoires than did primary tumors (Figure 6E), consistent with our data above showing that greater TCR evenness in tumors correlates with fewer tumor-distinct clones (Supplemental Figure 8E). In stark contrast though, the three metastases we were able to analyze from our lung cohort harbored more tumor-distinct clones than any other metastases in our study and lower evenness (Figure 6E, F). Together these data support the concept that lung-cohort patients are uniquely able to generate ongoing, clonal T cell responses to critically immunogenic antigens in both their primary and metastatic tumors, likely contributing to greater peripheral TCR diversity and possibly suppression of liver metastases.

## Discussion

Previous efforts to dissect PDAC tumors into subtypes used unbiased approaches with gene expression, proteomics, or metabolomics data to describe mutually exclusive subsets and defining signatures^25,26,28,29,31,33^. We approached the problem differently by classifying tumors based on a well-known clinical outcome associated with metastatic organotropism; specifically, longer-term survival among patients with lung-avid, liver-averse disease and poor outcomes with liver-avid disease^14–16,63^. We generated two new large datasets obtained from only patients with proven adenocarcinoma in the pancreas or Ampulla of Vater tumors that were histologically pancreaticobiliary: 1) a 290 specimen RNA-seq dataset with genomic alterations from tumor-enriched regions from primary and metastatic tumors and 2) TCRB sequencing of 289 blood and 175 matched tumor samples, mostly overlapping with the RNA-seq dataset. As previously reported^24^, tumors from patients in the lung cohort were unlikely to be categorized as the basal-like subtype; whereas classical subtype tumors were common in both cohorts. Uniquely, we found that patients with classical-subtype primary tumors fared significantly worse if their disease was liver-avid rather than lung-avid and liver-averse. This finding led us to investigate whether a unique gene signature exists for liver-avid tumors apart from gene-expression related to previously described subtypes.

We extracted a gene-expression signature for liver-avid primary tumors (pORG) that was devoid of genes overexpressed in basal-like primary tumors, addressing the overlap between these two subtypes and demonstrated that the pORG signature can independently predict patient outcome. High pORG and liver-avid primary tumors were enriched for pathways indicating ongoing replication stress and the response to replication stress by engagement of DNA repair processes, along with an increase in the activity of cell cycle drivers and mitogenic signaling (Figure 3 and 4, Supplemental Figure 6). Liver-avid tumors with somatic alterations in DDR genes had some of the highest pORG signature scores (Figure 3F), suggesting that PDAC tumor cells can avoid the detrimental effects of ongoing DNA damage by adopting strong RS response mechanisms. Conversely, patients live longer if their PDAC tumors have low pORG scores (Figure 2C), particularly if they harbor a DDR gene mutation (Figure 3E) and likely fail to productively respond to replication stress caused by DDR defects. A high pORG signature and ongoing response to replication stress was also associated with chronic interferon signaling, which unlike an acute type 1 interferon response, can induce an IFN-related DNA Damage Resistance Signature (IRDS)^53–55^ and reduced tumor immunity^56^.

Diverse TCRB repertoires associate with positive outcomes in PDAC patients^41,64^. Consistent with these reports, we identified high TCR evenness/diversity in peripheral blood from lung-cohort patients relative to patients in the liver cohort and most other patients with PDAC (Figure 5C and Supplemental Figure 7C) suggesting that there is something unique about tumor immunity in patients with lung-avid, liver-averse disease. We also found that lung-cohort patients had more T cell clones found in their tumors that were absent in paired blood samples compared to liver-cohort patients (Figure 5D); suggesting that there are new clonal responses in lung-cohort tumors that have not yet emigrated to blood, consistent with greater diversity building in the periphery over time (Supplemental Figure 7D). There are several possible explanations for enhanced T cell clonal development in lung-cohort tumors, including: a predilection for diversity in the pre-immune repertoire, greater adjuvanticity of the tumor (including the microenvironment) that promotes responses to many antigens, or a greater number of immunogenic neoepitopes or other tumor associated antigens generated in the tumor over time. A thorough comparison of immune competence in lung-cohort patients along with the immune contexture and neoepitope burden of their tumors may help parse these etiologies. It is also possible that only patients who stochastically develop a diverse T cell repertoire directed toward the right antigens end up with liver-averse disease.

Our results suggest that patients who develop liver metastases (a common outcome in PDAC) – relative to those with lung-avid, liver-averse disease – are more likely to generate clonal responses against persistent neoepitopes present since tumor initiation, at the expense of generating clonal responses to critically immunogenic tumor-associated antigens as they arise over time. This is consistent with a liver-avid primary tumor environment that favors weakly clonal responses of T cells prone to exhaustion over robust naïve T cell expansion against newly arising antigens^61^. Tumors are regularly refreshed with new T cell clones from the blood^65^, which in liver-cohort tumors may lead to a repeated cycle of T cell tumor exit, tumor entry, and weak clonal responses to a small pool of tumor initiating antigens. Our data supports that the distinction between high frequency T cell clones in liver-cohort and lung-cohort tumors (Figures 6A and C, respectively) is that the former represents clonal responses to a small pool of tumor initiating antigens that are decreasingly effective and the latter represents T cell expansion against new immunogenic antigens on which many lung-cohort patients have converged. In addition to this, we found that lung-cohort patients were uniquely able to generate tumor-distinct clones in both primary and metastatic tumors (Figure 6F), indicating that ongoing T cell clonal responses at several tumor sites may contribute to systemic resistance to PDAC progression.

In this report we identified a gene-expression signature representing ongoing replication stress response (i.e., high DNA replication and persistent mitogenesis and metabolism, along with concurrent DNA damage repair processes) that is associated with liver-avid, primary PDAC tumors. Tumors with low enrichment for this signature were less aggressive, particularly if DDR-participating genes were altered, suggesting that low pORG tumors may be more sensitive to therapeutics that interrupt DDR pathways. In concert with the low RS response of low pORG tumors, we found a high density of T cells and TCR sequences (Figure 4F and 5A) – one factor that may contribute to better outcomes for patients with low pORG tumors. A small subset of patients with low pORG tumors may direct those dense T cell infiltrates toward ongoing, clonal responses to critically immunogenic antigens leading to PDAC-relevant TCR diversity in the periphery and liver-averse disease. Our results point to two avenues for therapeutic intervention that may function additively: 1) exploitation of tumors lacking RS response mechanisms by promoting DNA damage (e.g., platinum chemotherapeutics) and interrupting DDR pathways (e.g., PARP inhibitors), and 2) immunomodulatory approaches (e.g., vaccines and immune checkpoint inhibition) that promote ongoing T cell responses to new tumor-associated antigens over time.

## Acknowledgments

We thank Jen Jen Yeh and laboratory for advice in performing PurIST analyses. We thank members of the Brenden-Colson Center for Pancreatic Care, the Sears Lab, and the Brody Lab for helpful discussions. We thank the Oregon Pancreas Tissue Registry, the OHSU Research Data Warehouse, and the Knight Biomedical Library for providing de-identified patient metadata and specimens, and the Knight Biostatistics Shared Resource for helpful discussion. The Knight shared resources are supported by Knight Cancer Institute P30 CA69533. These studies were supported by NIH-NCI U54CA209988, U01CA224012 and R01s CA186241, CA196228 to RCS; NIH-NCI 1U01 CA253472 and 5U01 CA217842 to GBM; and NIH-NCI R01 CA212600, U01CA224012-03, AACR Grant-15-90-25-BROD, and the Hirshberg Foundation to JBM; and foundation support from the Brenden-Colson Center for Pancreatic Care.

