## Supporting Information for "Ongoing Replication Stress Response and New Clonal T Cell Development Discriminate Between Liver and Lung Recurrence Sites and Patient Outcomes in Pancreatic Ductal Adenocarcinoma"

### Supplemental Figure 1

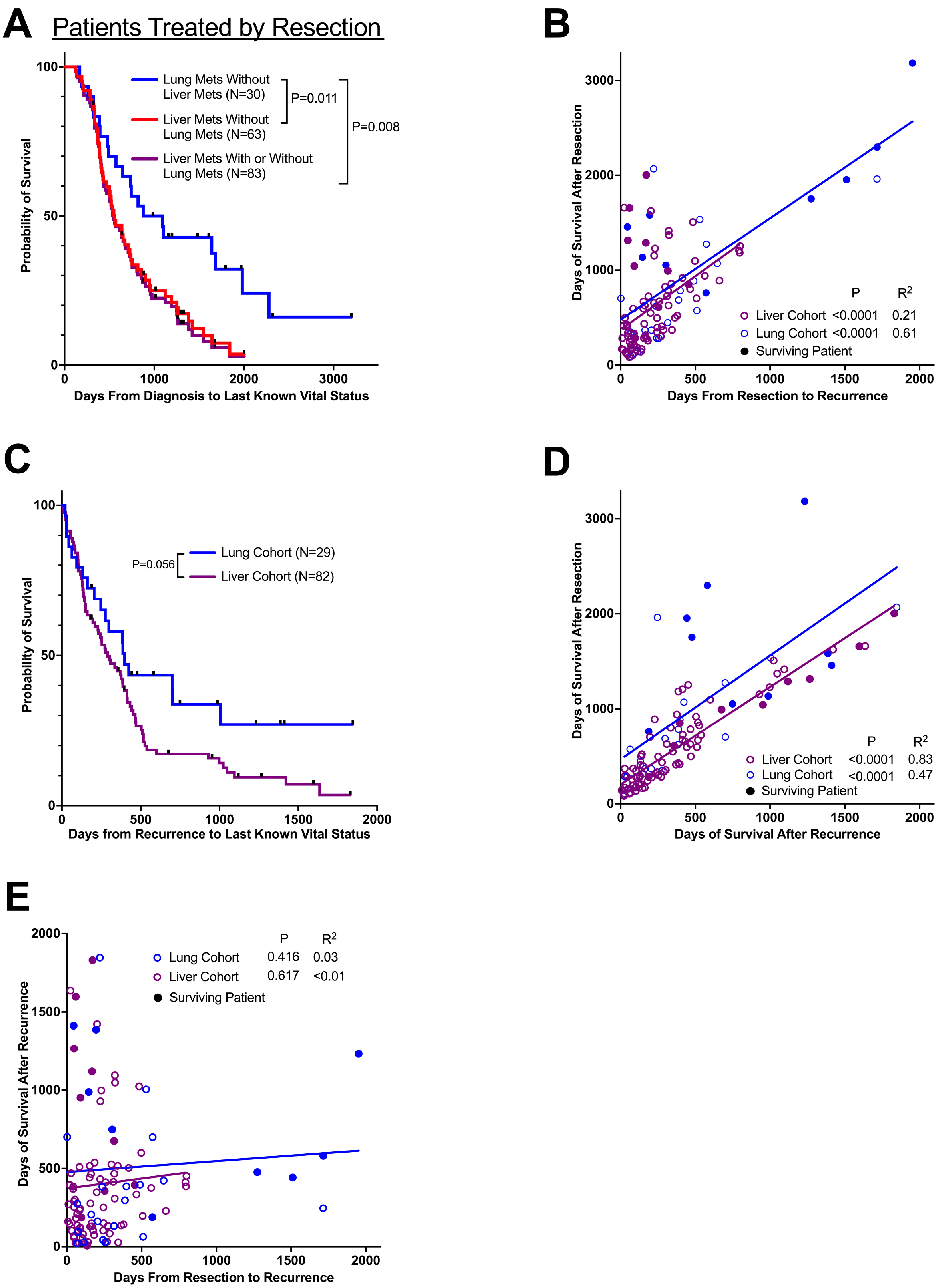


Supplemental Figure 1: A) Overall survival for patients treated by surgical resection of primary tumors. B) Pearson correlation between days to recurrence and survival after resection. C) Survival after recurrence D) Pearson correlation between survival after recurrence and survival after resection. E) Pearson correlation between time to recurrence after resection and survival after recurrence. For charts with survival endpoints, patients not deceased were censored at last known vital status. Patients who died <30 days after resection were omitted. A Mantel-Cox test was used to compare all Kaplan-Meier survival and recurrence curves.

### Supplemental Figure 2

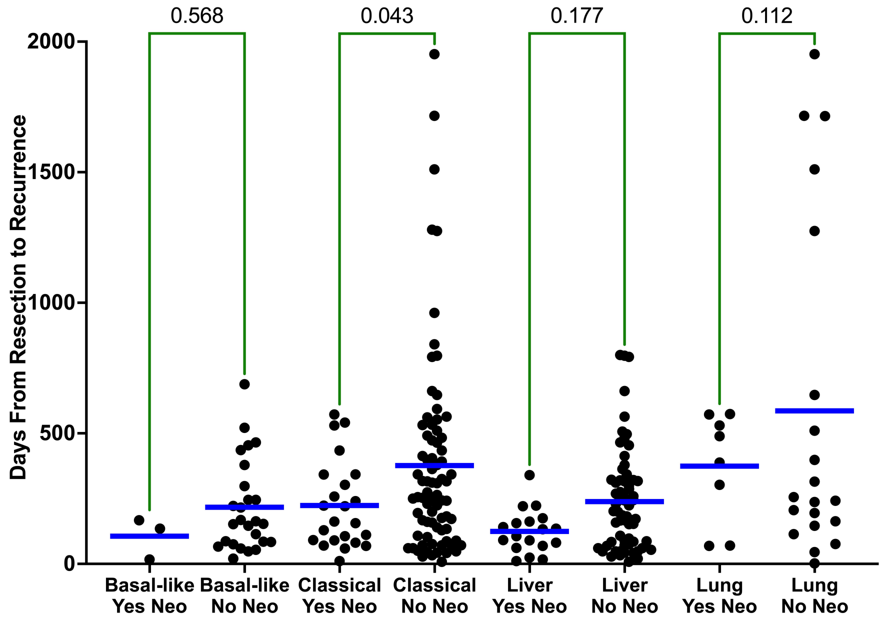


Supplemental Figure 2: The influence of neoadjuvant treatment (Neo) on time to recurrence after resection of primary tumors for patients in the liver cohort (Liver) and lung cohort (Lung) or categorized by PurIST tumor subtype. One-way ANOVA was used to calculate indicated P values. For the two patients with more than one specimen analyzed, the resected primary tumor was used for subtype assignment. Blue bars represent means.

### Supplemental Figure 3

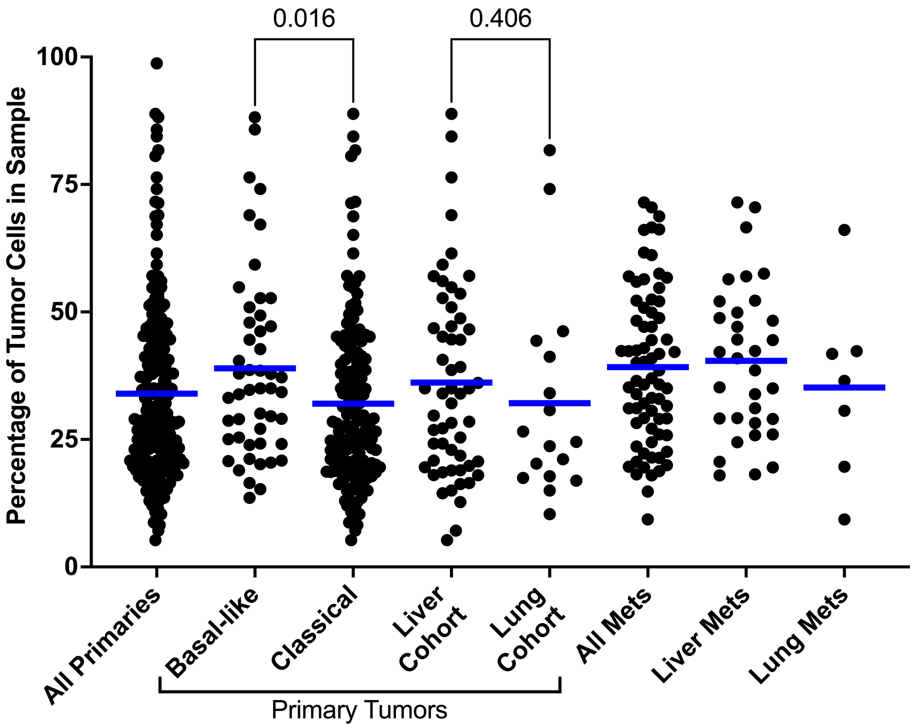


Supplemental Figure 3: The percentage of tumor cells in samples analyzed by RNASeq determined by mutant allele frequencies from the Tempus xT genomic alteration panel. One-way ANOVA was used to calculate indicated P values. For the two patients with more than one specimen analyzed, the resected primary tumor was used for subtype assignment. Blue bars represent means.

### Supplemental Figure 4

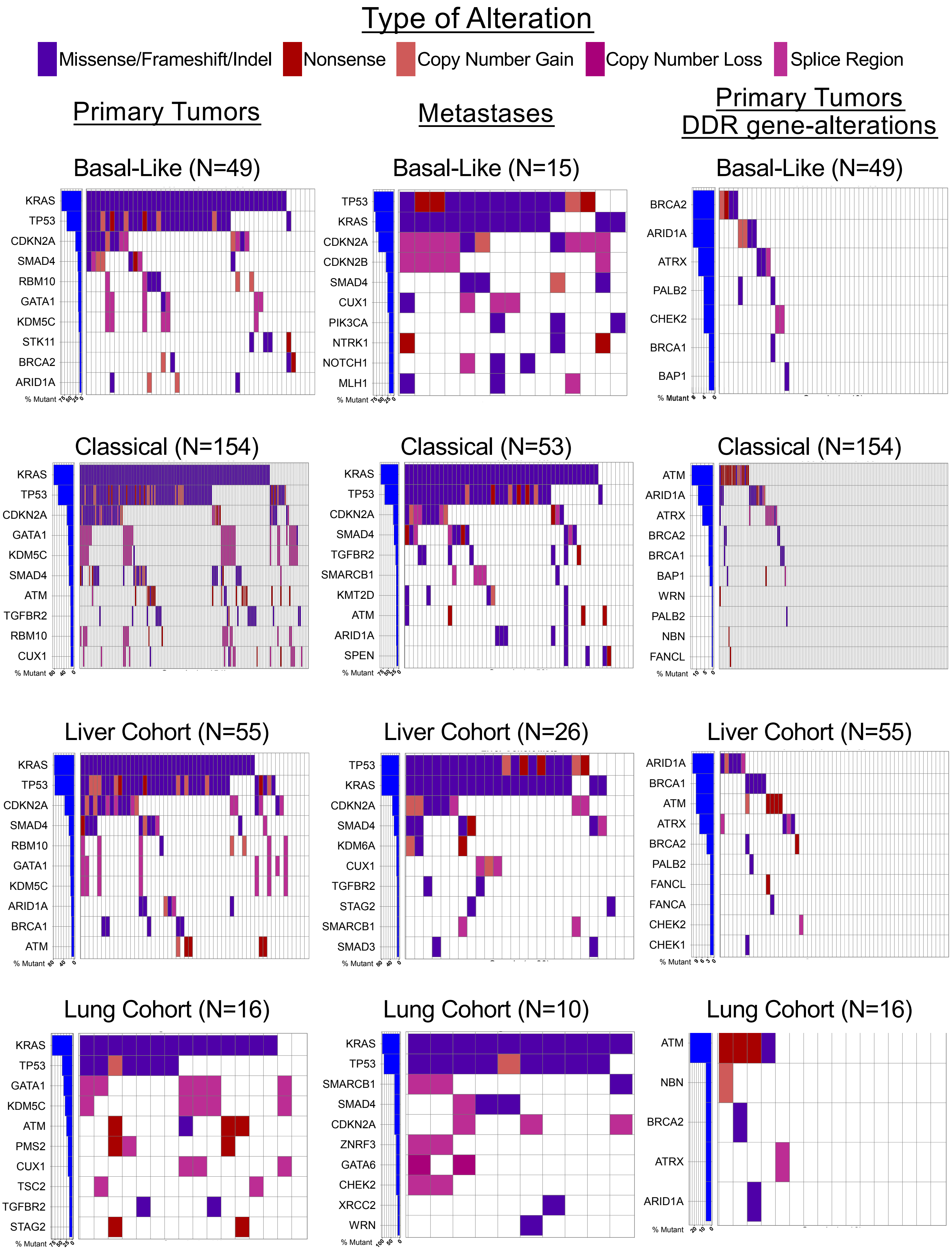

Supplemental Figure 4: The top ten most frequently altered genes and their alteration types are displayed by cohort for metastatic tumors (left plots), primary tumors (middle plots), and for all DDR-related genes among primary tumors (right plots). DDR genes are not included on the plot if no samples in the cohort had an alteration. Blue bars to the left of each plot represent the percentage of tumors with an alteration in each gene.

### Supplemental Figure 5

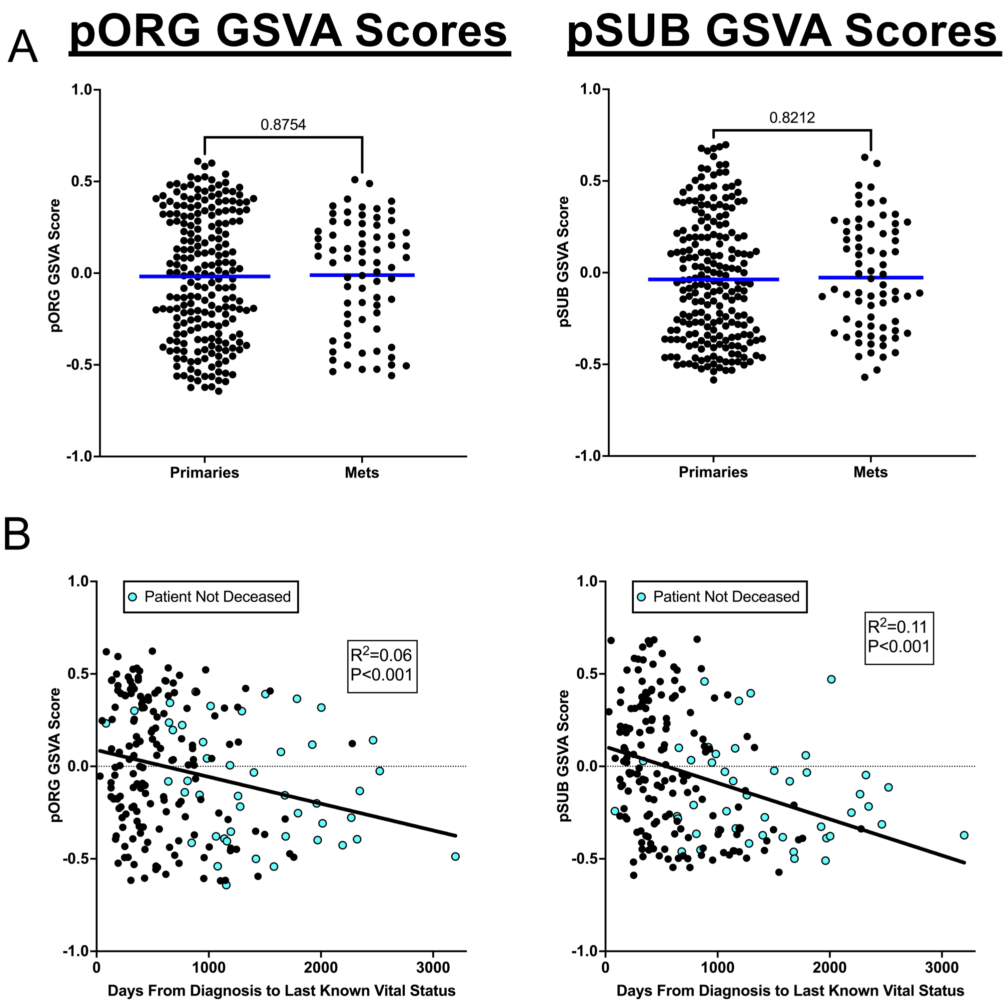


Supplemental Figure 5: A) GSVA scores for the pORG (left panels) and pSUB (right panels) are shown for primary and metastatic tumors. B) Pearson correlation between pORG (left panel) and pSUB GSVA (right panel) scores and overall survival. Patients who died within 30 days after resection are not shown.

### Supplemental Figure 6

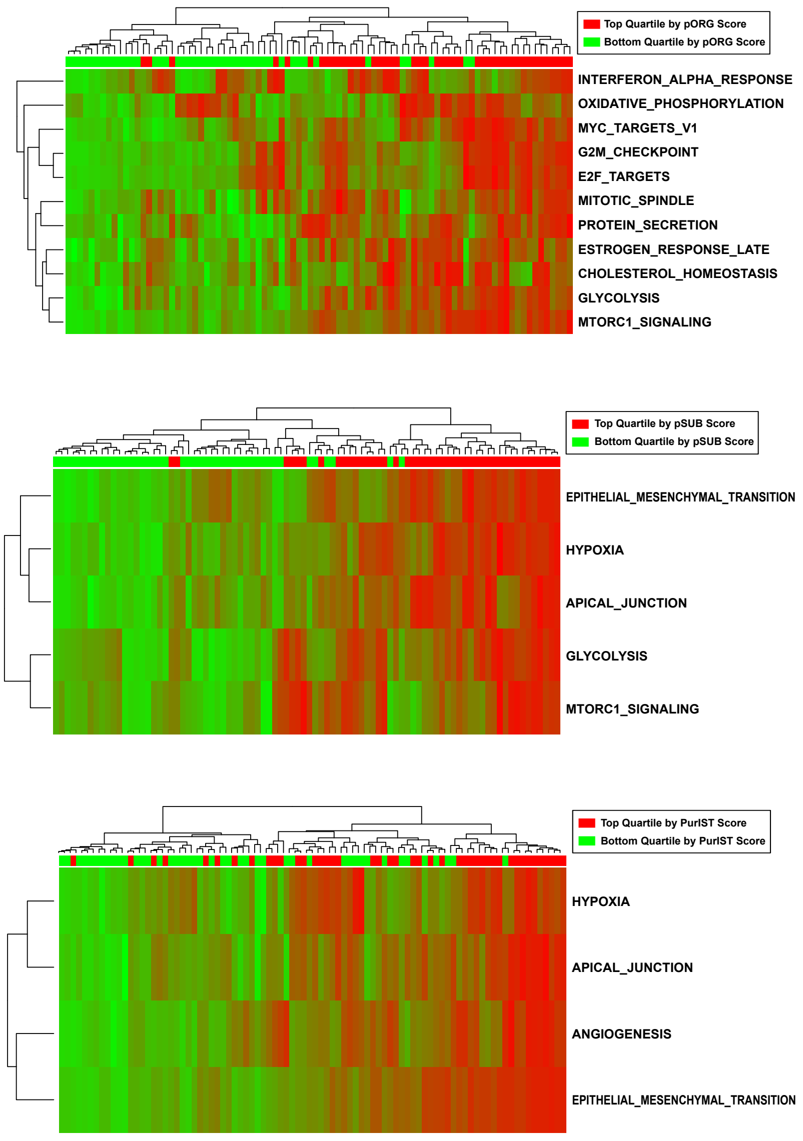


Supplemental Figure 6: Heatmaps of GSVA scores (high=red, low=green) for primary tumors from the TCGA PAAD dataset showing pathway (NES >1.7 and nominal P<0.05) and specimen clustering from the cohorts indicated on each plot. Shown are the same pathways displayed in Figure 4.

### Supplemental Figure 7

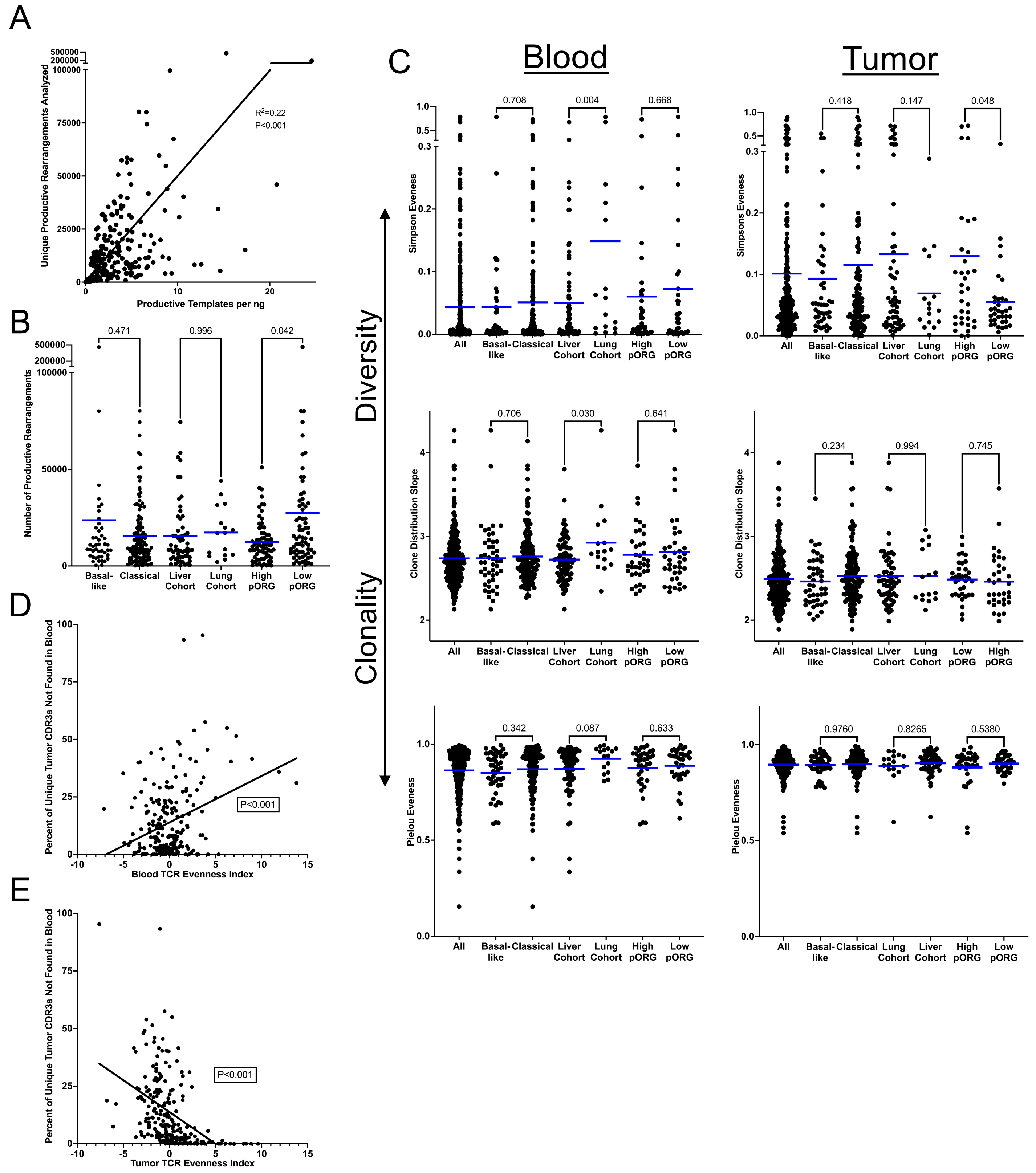


Supplemental Figure 7: A) Pearson correlation between productive templates per ng of tumor tissue and unique, productive rearrangements analyzed for all tumors. B) The total number of unique productive TCR CDR3β rearrangements analyzed for the indicated cohorts. C) Shown are three different metrics of the evenness of TCRB CDR3β repertoires from blood (left panels) and tumor (right panels) samples in the indicated cohorts. Blue bars represent means. D) Pearson correlation between the percentage of tumor-distinct clones and blood TCR CDR3β Evenness Index scores. E) Pearson correlation between the percentage of tumor-distinct clones and blood TCR CDR3β Evenness Index scores.

### Supplemental Table 1

A

#### Primary Tumors

| Gene Altered | Percent Altered in Liver Cohort | Percent Altered in Lung Cohort | Fisher's Test P Value |
|---|---|---|---|
| PMS2 | 2 | 19 | 0.034 |
| PTEN | 0 | 13 | 0.048 |
| TP53 | 75 | 44 | 0.033 |

| Gene Altered | Percent Altered in Basal-like | Percent Altered in Classical | Fisher's Test P Value |
|---|---|---|---|
| ABL1 | 6 | 1 | 0.044 |
| ATM | 0 | 13 | 0.005 |
| CUX1 | 2 | 12 | 0.048 |
| GNAS | 0 | 10 | 0.024 |
| MKI67 | 6 | 0 | 0.013 |
| SMARCA4 | 6 | 1 | 0.044 |
| TGFBR2 | 2 | 12 | 0.050 |

B

#### Metastases

| Gene Altered | Percent Altered in Basal-like | Percent Altered in Classical | Fisher's Test P Value |
|---|---|---|---|
| CDKN2A | 67 | 25 | 0.004 |
| CDKN2B | 33 | 2 | 0.001 |
| CUX1 | 27 | 6 | 0.038 |
| EBF1 | 13 | 0 | 0.046 |
| FANCD2 | 13 | 0 | 0.046 |
| MLH1 | 20 | 0 | 0.009 |
| MTAP | 40 | 8 | 0.006 |
| NOTCH1 | 20 | 0 | 0.009 |
| PIK3CA | 20 | 2 | 0.031 |
| TRAF3 | 13 | 0 | 0.046 |

Supplemental Table 1: Shown are all genes significantly altered (Two-tailed Fishers exact test) between the indicated cohorts and the frequency of samples with the gene altered in each cohort for A) primary tumors and B) metastatic tumors.

### Supplemental Table 2

| Patient | Specimen | PurIST Subtype | PurIST Detailed Subtype | PurIST Score | Liver Cohort | Lung Cohort | Pair Type | KRAS | TP53 | CDKN2A | SMAD4 |
|---|---|---|---|---|---|---|---|---|---|---|---|
| ST-00007307 | ST-00007307-T | classical | strong classical | 0.00549982 | | YES | Primary/Met | G12V | | | |
| ST-00007307 | ST-00007307-M | classical | strong classical | 0.00549982 | | YES | Primary/Met | G12V | | CNL | |
| ST-00006291 | ST-00006291-T | classical | strong classical | 0.01380528 | YES | | Primary/Met | | | CNL | |
| ST-00006291 | ST-00006291-M | classical | likely classical | 0.13368768 | YES | | Primary/Met | | | | |
| ST-00010984 | ST-00010984-T | classical | strong classical | 0.0089058 | | | Primary/Met | G12C | Splice | | |
| ST-00010984 | ST-00010984-M | classical | likely classical | 0.30353193 | | | Primary/Met | G12C | Splice | | |
| ST-00014524 | ST-00014524-A-T | classical | likely classical | 0.32504852 | YES | | Primary/Met | | Splice | CNL | |
| ST-00014524 | ST-00014524-M | basal-like | strong basal-like | 0.90222402 | YES | | Primary/Met | | Splice | CNL | |
| ST-00015839 | ST-00015839-T | classical | strong classical | 0.00109571 | | YES | Primary/Met | Q61H | R273H | | |
| ST-00015839 | ST-00015839-M | classical | strong classical | 0.00109571 | | YES | Primary/Met | Q61H | R273H | CNL | CNL |
| ST-00017804 | ST-00017804-T | classical | strong classical | 0.00549982 | | YES | Primary/Met | Q61R | R175H | | |
| ST-00017804 | ST-00017804-M | classical | strong classical | 0.00109571 | | YES | Primary/Met | Q61R | R175H | CNL | |
| ST-00017838 | ST-00017838-T | basal-like | strong basal-like | 0.97822823 | YES | | Primary/Met | G12V | R337L | | CNL |
| ST-00017838 | ST-00017838-M | classical | likely classical | 0.33333579 | YES | | Primary/Met | G12V | R337L | | |
| ST-00017945 | ST-00017945-T | basal-like | likely basal-like | 0.78592507 | | | Primary/Primary | G12D | R196* | | |
| ST-00017945 | ST-00017945-T2 | basal-like | lean basal-like | 0.56008026 | | | Primary/Primary | G12D | R196* | | |
| ST-00004898 | ST-00004898-T | classical | strong classical | 0.02525726 | | | Primary/Met | G12V | K292* | | |
| ST-00004898 | ST-00004898-M | classical | strong classical | 0.00549982 | | | Primary/Met | G12V | K292* | | V492D |
| ST-00019598 | ST-00019598-T | classical | strong classical | 0.0404529 | | | Primary/Primary | G12D | | | |
| ST-00019598 | ST-00019598-T2 | classical | likely classical | 0.16494877 | | | Primary/Primary | | | | |
| ST-00019429 | ST-00019429-T | classical | strong classical | 0.00109571 | | | Primary/Met | No Data | | | |
| ST-00019429 | ST-00019429-M | classical | strong classical | 0.00109571 | | | Primary/Met | | | | |
| ST-00019601 | ST-00019601-T | classical | strong classical | 0.01370931 | | | Primary/Met | G12V | L257Q | | |
| ST-00019601 | ST-00019601-M | classical | strong classical | 0.06155242 | | | Primary/Met | G12V | | M52fs | |

Supplemental Table 2: All paired specimens (from the same patient) are shown along with their PurIST, liver, and lung cohort assignments and genomic alterations in the four most frequently altered genes in PDAC.
